# Nuclear speckles regulate HIF-2α programs and correlate with patient survival in kidney cancer

**DOI:** 10.1101/2023.09.14.557228

**Authors:** Katherine A. Alexander, Ruofan Yu, Nicolas Skuli, Nathan J. Coffey, Son Nguyen, Christine Faunce, Hua Huang, Ian P. Dardani, Austin L. Good, Joan Lim, Catherine Li, Nicholas Biddle, Eric F. Joyce, Arjun Raj, Daniel Lee, Brian Keith, M. Celeste Simon, Shelley L. Berger

## Abstract

Nuclear speckles are membrane-less bodies within the cell nucleus enriched in RNA biogenesis, processing, and export factors. In this study we investigated speckle phenotype variation in human cancer, finding a reproducible speckle signature, based on RNA expression of speckle-resident proteins, across >20 cancer types. Of these, clear cell renal cell carcinoma (ccRCC) exhibited a clear correlation between the presence of this speckle expression signature, imaging-based speckle phenotype, and clinical outcomes. ccRCC is typified by hyperactivation of the HIF-2α transcription factor, and we demonstrate here that HIF-2α drives physical association of a select subset of its target genes with nuclear speckles. Disruption of HIF-2α-driven speckle association via deletion of its <u>s</u>peckle targeting <u>m</u>otifs (STMs)—defined in this study—led to defective induction of speckle-associating HIF-2α target genes without impacting non-speckle-associating HIF-2α target genes. We further identify the RNA export complex, TREX, as being specifically altered in speckle signature, and knockdown of key TREX component, ALYREF, also compromises speckle-associated gene expression. By integrating tissue culture functional studies with tumor genomic and imaging analysis, we show that HIF-2α gene regulatory programs are impacted by specific manipulation of speckle phenotype and by abrogation of speckle targeting abilities of HIF-2α. These findings suggest that, in ccRCC, a key biological function of nuclear speckles is to modulate expression of a specific subset of HIF-2α-regulated target genes that, in turn, influence patient outcomes. We also identify STMs in other transcription factors, suggesting that DNA-speckle targeting may be a general mechanism of gene regulation.

**HIGHLIGHTS:**

– Nuclear speckles shown to reproducibly vary in cancer, predicting patient survival in ccRCC
– HIF-2α drives DNA/gene-speckle contacts dependent on identified speckle targeting motifs within HIF-2α
– Putative speckle targeting motifs are highly enriched among regulators of gene expression
– Partitioning of transcription factor functional programs may be a major biological function of nuclear speckles

## INTRODUCTION

Nuclear speckles are dynamic nuclear bodies characterized by high local concentrations of RNA binding proteins and specific non-coding RNAs (reviewed in ^1–3^). Previously known as “interchromatin granule clusters” and “splicing speckles”, nuclear speckles occupy regions reduced in DNA content and take up ∼10-30% of the nuclear volume. Nuclear speckles are defined in immunofluorescence experiments by speckle markers RBM25, SON, and/or SRRM2, of which SON and SRRM2 are together responsible for nuclear speckle structural integrity^4,5^. Biochemical fractionation, affinity labelling, and imaging methods show that nuclear speckles contain numerous RNA binding proteins and gene expression regulators involved in chromatin organization, transcription, splicing, polyadenylation, RNA modification, and RNA nuclear export^6–8^. While it has been noted that speckles vary across cell types and change in response to cellular perturbations^3,9–12^, systematic quantitation of speckle variation and evaluation of the consequences of such differences have been historically challenging given the complexity of speckle properties and contents.

Although the contents of speckles suggest multifaceted roles in regulating chromatin dynamics and gene expression (reviewed in ^13^), the overarching biological function of nuclear speckles remains enigmatic. In support of gene-activating roles, the most highly active genes tend to be adjacent to speckles^14,15^. Specific genes also change their position relative to nuclear speckles during cellular stress (heat shock^16–20^ and p53 activation^21^) and differentiation (erythropoiesis^22^). In the context of p53 activation, not all p53 target genes alter their speckle association status, however target genes that experience p53-mediated increases in speckle association display higher induction compared to target genes that lack increased speckle association^21^. Thus, rather than acting as an on/off switch, speckles may amplify expression of a select subclass of transcription factor target genes, conferring an expression advantage of specific gene sets. Altogether, these studies support that speckles occupy a distinct gene regulatory niche that is particularly important in changing environments such as stress responses or the process of cell differentiation.

Gene regulation by nuclear speckles can theoretically be accomplished by altering which genes are positioned nearby speckles and/or by changes in nuclear speckles themselves. Both mechanisms are evident in viral exploitation of speckle pathways (reviewed in ^13^). Mutations and altered expression of speckle-resident proteins are also evident in cancer, and speckle-resident protein mutations are recurrently found in developmental disorders, described as “speckleopathies” (reviewed in ^13,23^). While links between speckle-resident proteins and human pathologies are compelling, the extent and underlying logic of nuclear speckle variability in human health and disease, and gene expression repercussions of such variation, are unclear.

We reported previously that DNA-speckle association is driven by the transcription factor p53^21^. Given the existence of regulated DNA-speckle association during heat shock and erythropoiesis^16,17,20,22^, processes involving other specific transcription factors (HSF1 and GATA1, respectively), we hypothesize that speckle targeting could be a broadly-utilized transcription factor gene regulatory mechanism in both normal and pathological contexts. However, whether other transcription factors drive DNA-speckle association is unknown and we currently lack methods to predict which transcription factors have this ability.

Here, we investigate speckle phenotypes that correlate with cancer and discover two alternative broad signatures of gene expression of speckle resident proteins. We find that one signature correlates with poor patient survival specifically in clear cell renal cell carcinoma (ccRCC). ccRCC is typically driven by hyperactivation of the HIF-2α transcription factor, and we show HIF-2α mediates DNA-speckle association— leading to extensive mechanistic insight into how transcription factors regulate target gene association with speckles. We integrate tissue-culture-based functional approaches with genomic and imaging analyses in ccRCC tumors, providing strong support for speckles as a critical gene regulatory layer. Our findings thus connect nuclear speckle-based gene regulation to cancer patient outcomes, providing a new link between nuclear architecture and human disease.

## RESULTS

### A recurring speckle protein gene expression signature predicts patient outcomes in clear cell renal cell carcinoma (ccRCC)

Although speckle-resident proteins are mutated in cancers and developmental disorders (reviewed in ^13,23^), methods to systematically evaluate nuclear speckle phenotypes in altered states are lacking. We undertook characterization of nuclear speckle variation in human cancer, utilizing RNA expression of genes encoding speckle-resident proteins as a proxy for speckle phenotypes. We extracted 446 speckle-resident proteins based on speckle-localization annotations from the Human Protein Atlas^8^ (**Fig 1Ai**) and estimated speckle phenotypes from their RNA expression in The Cancer Genome Atlas (TCGA) using Principal Component Analysis (PCA)(**Fig 1Aii**; see **Fig S1A** for detailed schematic). Comparing speckle protein gene expression contributions to patient variation between cancer types (derived from PCA analysis; see Methods for detailed explanation), we observed remarkable correlations between cancer types (**Fig 1Aiii**, strong correlations are orange and red), indicating that speckle protein gene expression varies reproducibly in cancer.

**Figure 1.**
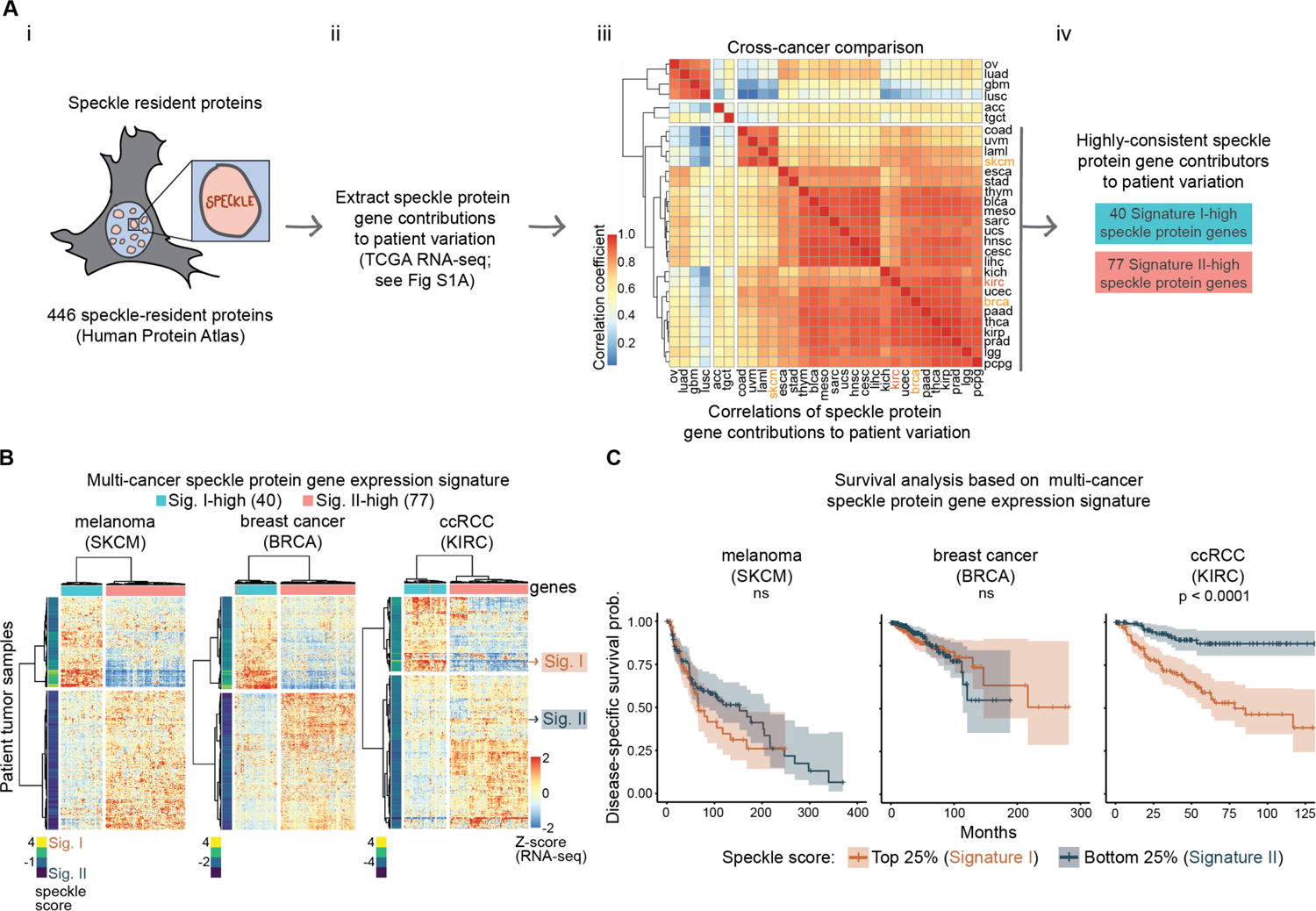
A recurring speckle signature predicts patient outcomes in ccRCC. A) Schematic showing generation of multi-cancer speckle signature. Proteins residing within speckles were identified (i), their expression evaluated (ii), contributions to patient variation compared (iii; Pearson’s correlations), and consistent speckle protein gene contributors to patient variation were identified (iv). See also **Fig S1A**. B) Heatmaps showing z-scores of speckle signature speckle protein gene RNA expression in melanoma (SKCM), breast cancer (BRCA), and renal cell carcinoma (KIRC). Bar on left represents speckle scores. Bar above represents Signature I-(blue) or Signature II-(pink) high speckle protein genes. C) Kaplan Meier plots separating cancer cohorts by the top and bottom 25% of speckle scores. See also **Fig S1B** and **Table S1**.

Based on this consistent speckle protein gene expression variation across many cancer types, we generated a multi-cancer 117 gene “speckle signature” containing speckle protein genes that consistently contributed to patient variation (**Fig 1Aiv**; see Methods and **Fig S1A** for how speckle protein genes were selected). This included 40 “Signature I-high” speckle protein genes and 77 “Signature II-high” speckle protein genes that were consistently reciprocally expressed from one another, and that separated tumor samples into two groups based on their expression (**Fig 1B**). We assigned each patient a speckle signature score based on the collective expression of these 117 speckle protein genes (**Fig 1B**, speckle score to the left of each heatmap; see Methods) and used this quantitative measure for Kaplan-Meier analysis of disease outcomes.

We assessed overall and disease-specific survival, separating patients by the top versus bottom quartiles or positive versus negative speckle scores (4 total analyses; **Fig S1B**). Of 24 cancers with consistent speckle protein gene contributions to patient variation (right grey bar in **Fig 1Aiii**), 21 showed no correlation between speckle signature and patient outcomes for any survival measurement, as shown by examples of melanoma (SKCM) and breast cancer (BRCA) (**Fig 1C**, left panels; **Table S1**). In head and neck cancer (HNSC), speckle Signature I was slightly correlated with poor overall survival in the quartile analysis (**Table S1**), while in ovarian cancer (OV), speckle Signature II slightly correlated with poor outcomes in the quartile analysis (**Fig S1B**, top left).

Contrasting with other cancer types, speckle signature in clear cell renal cell carcinoma (ccRCC; called KIRC in TCGA data) showed remarkable correlation with patient outcomes (**Fig 1C**, right; **Fig S1B** bottom; **Table S1**). In ccRCC, patients expressing speckle Signature I had significantly worse outcomes compared to patients with speckle Signature II. Further, although speckle Signature I was more prevalent in later stage ccRCC (**Fig S1C**; bottom bars), speckle Signature I also occurred in early stage ccRCC (**Fig S1C**; top bars) and was predictive of survival in both early and late stages (**Fig S1D**). Hence, speckle signature can be an early-occurring predictor of patient outcomes in ccRCC. Of note, normal adjacent tissues of ccRCC, and most other cancer types, showed speckle Signature II expression patterns (**Fig S1E**), indicating that speckle Signature I reflects a dysfunction that emerges in tumors. Overall, these findings demonstrate that while the two speckle signatures were evident in 24 cancer types, they correlate strongly with patient outcomes specifically in ccRCC.

### A putative speckle targeting motif (STM) is shared between HIF-2α and p53 and re-occurs among regulators of gene expression

We next sought to illuminate why patient survival in ccRCC was particularly correlated with speckle signature differences. One feature of ccRCC that distinguishes it from other cancer types is that ccRCC is typified by inactivating mutations in the *VHL* gene (reviewed in ^24^), whereas other cancers within the TCGA PanCan dataset have more heterogeneous etiologies. The pVHL protein encoded by *VHL* is a negative regulator of hypoxia-inducible transcription factors HIF-1α and HIF-2α, of which HIF-2α is proposed to be the oncogenic driver^24–26^. Given our previous demonstration of DNA-speckle targeting abilities by the p53 transcription factor, we hypothesized that HIF-2α may be a second example of a stress-responsive transcription factor that regulates association between specific genes and nuclear speckles.

HIF-2α and p53 belong to different transcription factor classes^27–29^. However, using EMBOSS Matcher^30^ to perform local alignment, we discovered a 29 amino acid region as the best match between p53 and HIF-2α, with 55% sequence similarity (**Fig S2A**). This region comprised p53 amino acids 62-90, and encompasses the exact region required for DNA/target gene association with speckles that we mapped previously (p53 amino acids 62-77)^21^. The aligned sequence is characterized by prolines spaced every 5 amino acids (**Fig 2A**, labelled with *), with one proline preceded by a threonine (**Fig 2A**, green highlight). Examination of HIF-2α revealed a second region of spaced prolines with a serine/proline (SP) dinucleotide in the same relative position, and a similar sequence was present in GATA1 (**Fig 2A**), a transcription factor active during erythropoiesis, which also involves regulated DNA-speckle association^22,31^. Based on these findings, we termed this protein region the putative <u>s</u>peckle targeting <u>m</u>otif (STM). Pairwise Clustal Omega alignment between HIF-2α and HIF-1α showed that HIF-1α lacks the proline to the N-terminus of the TP dinucleotide found in the first HIF-2α STM (**Fig S2B**; **Fig S2C**, top red circle), and thus does not have prolines spaced every 5 amino acids, whereas the second HIF-2α STM is absent in HIF-1α (**Fig S2B** and **Fig S2C**). Hence, putative STMs are found within p53, HIF-2α, and GATA1, but are divergent between HIF-2α and HIF-1α.

**Figure 2.**
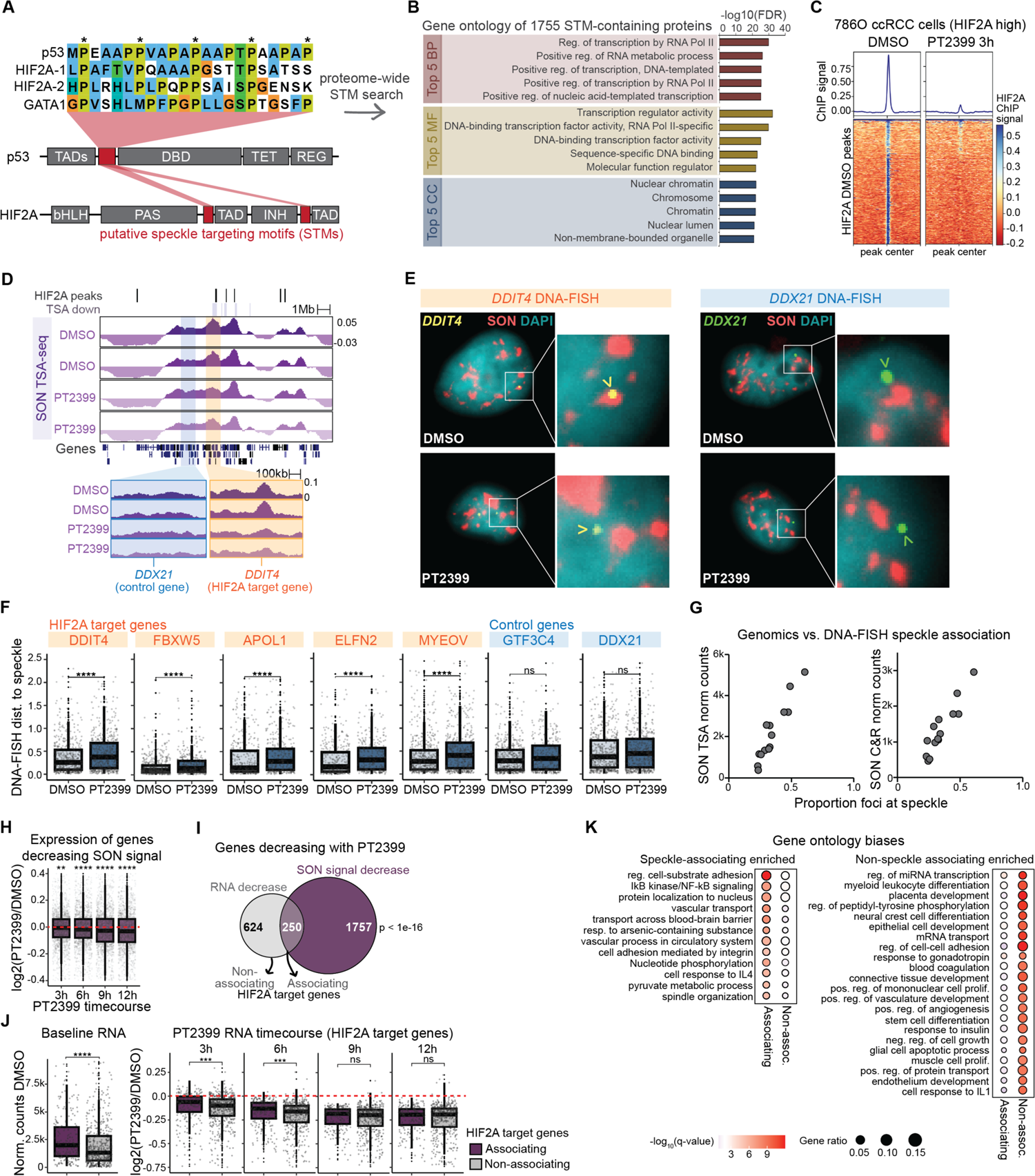
HIF-2α maintains DNA-speckle contacts of its target genes in 786-O ccRCC cells. A) Alignment of putative speckle targeting motifs (STMs) shared between p53, HIF-2α, and GATA1 with schematic of STM locations in p53 and HIF-2α. B) Top 5 Biological Processes (BP), Molecular Functions (MF), and Cellular Compartments (CC) found by Gene Ontology analysis of 1755 proteins containing putative STMs C) Metaplot and heatmap of HIF-2α ChIP-seq signal at HIF-2α binding peaks in 786-O cells treated with DMSO control (left) or HIF-2α inhibitor (PT2399, right). D) Genome browser tracks of SON TSA-seq in 786-O cells upon 3 hour DMSO or PT2399 treatment. Decreased SON regions (purple bars) and HIF-2α binding sites (black bars) are shown above tracks. *DDX21* (blue) and *DDIT4* (orange) regions with zoomed views were used for DNA-FISH in **E**. E) Example immunoDNA-FISH images of speckles (red; SON immunofluorescence), nuclei (cyan; DAPI), *DDIT4* DNA-FISH (yellow, left images), and *DDX21* DNA-FISH (green, right images). F) Quantification of immunoDNA-FISH loci distance to the nearest speckle (in µm). Each dot represents one locus. Significance calculated using Wilcoxon test. G) Relationship between SON TSA-seq (left) or SON Cut&Run (right) and proportion of foci at speckle measured by immunoDNA-FISH. Each dot represents one genomic locus. H) Expression change of PT2399 versus DMSO in a 786-O cell PT2399 timecourse. Each dot represents one gene decreasing SON signal. Significance calculated using T-test versus null hypothesis of no change. I) Venn diagram showing overlap of genes decreasing RNA expression and SON signal. Significance calculated using hypergeometric test. J) Baseline RNA expression (left), and RNA fold change in a PT2399 timecourse (right). Each dot represents one HIF-2α target gene that did (associating; purple box) or did not (non-associating; grey box) have speckle association decrease with PT2399. K) Gene ontology showing HIF-2α pathways biased toward associating (left) or non-associating (right) HIF-2α target genes. ** – p < 0.01; *** – p < 0.001; **** – p < 0.0001, ns – not significant For full list of STM-containing proteins and STM sequences, see **Table S2** and **Table S3**.

Using p53, HIF-2α, and GATA1 as instructive examples, we generated an algorithm for *de novo* identification of putative STM-containing proteins (see **Fig S2D** for criteria). In a proteome-wide search, we found 1755 proteins that contain putative STMs (**Table S2** – gene names; **Table S3** – STM sequences). Gene Ontology of STM-containing proteins revealed that they were profoundly enriched in transcription and gene regulation Biological Processes (BP), Molecular Functions (MF), and Cellular Compartments (CC) (**Fig 2B**).

591 STM-containing proteins were involved in “Regulation of gene expression” (591/5164 GO:0010468; FDR of enrichment < 1e-15), including >200 DNA-binding transcription factors (e.g. OCT4, KLF4, NOTO, RELA, MYCL, RUNX1). We also identified putative STMs among chromatin organizers, histone modifying enzymes, and proteins involved in enhancer activity, such as H3 Lys4 methyltransferases (e.g. KMT2A, KMT2B, KMT2C, and KMT2D), eleven members of the Mediator complex, and histone acetyltransferases (e.g. CPB, p300, KAT6A). STMs were also found within RNA binding and speckle-resident proteins themselves (e.g. SF3B1, CACTIN, SETD2, SRRM2 and SON; FDR of enrichment in GO:0016607 “nuclear speck” < 0.001) as well as several members of the nuclear pore complex (e.g. NUP98, TPR, NUP61). Hence, putative STMs re-occur among regulators of gene expression, spanning direct-DNA binding factors and beyond.

### HIF-2α maintains DNA-speckle contacts of its target genes in ccRCC cells

While hundreds of gene regulators contain STMs, it is unclear whether the presence of an STM predicts DNA-speckle targeting. To test speckle targeting functions of HIF-2α, we utilized 786-O ccRCC cells, a patient-derived cell line that has elevated endogenous HIF-2α^32^, and inhibited the interaction between HIF-2α and its obligate DNA-binding heterodimer ARNT/HIF-1β with the PT2399 small molecule^33,34^. ChIP-seq of HIF-2α with the inhibitor showed loss of HIF-2α genomic binding at 3 hours after PT2399 compared to vehicle treatment (**Fig 2C**). We used this time point to assess whether HIF-2α DNA binding is essential for speckle association, employing two genomic methods to measure DNA-speckle contacts. The first, SON TSA-seq, is a proximity-labeling approach that utilizes an antibody against the SON speckle marker protein to capture DNA that resides near speckles^14,19^. The second, SON Cut&Run, is a pA-MNase method that captures speckle-associated DNA by digesting chromatin fragments that surround SON^35^. We found high correlation between genomic SON signal from TSA-seq and Cut&Run (**Fig S2E**). SON TSA-seq and SON Cut&Run detected significant PT2399-dependent decreases in SON (padj < 0.01), and decreasing regions significantly overlapped between the two methods. However, each method detected decreases that were not detected by the other (**Fig S2F**). In comparing them, SON TSA-seq more effectively identified alterations at lower SON regions, while SON Cut&Run more readily detected changes at higher SON regions (**Fig S2G**). Distinct types of regions detected as differential between the two methods was not surprising given that they use distinct chemistries to identify speckle associated chromatin. In summary, we consider that both methods are valid, capturing different types of decreasing regions, we thus combined them in the analyses below.

As a third method to evaluate DNA-speckle contacts upon HIF-2α inhibition, we employed immunoDNA-FISH with immunofluorescence against SON to label speckles and DNA-FISH against gene loci to label DNA. We designed probe sets against regions that decreased in SON TSA-seq or Cut&Run signal upon PT2399 treatment (for example, **Fig 2D**, *DDIT4* orange box), as well as regions that did not change (for example, **Fig 2D**, *DDX21* blue box). We treated 786-O cells with DMSO control or PT2399, and found that HIF-2α inhibition resulted in the *DDIT4* HIF-2α target gene localizing farther from speckles, while the *DDX21* control gene did not change speckle association (**Fig 2E** – example images; **Fig 2F** – quantification). Likewise, *FBXW5*, *APOL1* and *ELFN2* HIF-2α target genes that significantly decreased SON in TSA-seq and Cut&Run showed reduced speckle association by immunoDNA-FISH, as did *MYEOV* (**Fig 2F**), which had decreased SON signal by Cut&Run, but not by TSA-seq. A second control gene, *GTF3C4*, which is within the same larger speckle domain as *FBXW5*, did not decrease SON signal in either genomic method, and did not show changes in speckle association by immunoDNA-FISH (**Fig 2F**). Comparing the immunoDNA-FISH-measured proportion of foci at speckles to the amount of SON signal for SON TSA-seq (**Fig 2G**, left) and SON Cut&Run (**Fig 2G**, right) for each gene, we observed similar correlations between imaging and genomic speckle-association measurements, supporting the contention that each method reflects the amount of DNA-speckle association. In total, these three orthogonal approaches demonstrate that inhibition of HIF-2α DNA binding lowers HIF-2α target gene speckle association, indicating a new mechanistic function of HIF-2α for maintaining DNA-speckle contacts in ccRCC cells.

### HIF-2α regulates speckle association of a functionally-distinct subset of target genes

Our findings indicate that, like p53, HIF-2α has a speckle targeting function that drives association between specific genes and nuclear speckles. Previously, we showed that p53 regulates speckle association of 20-30% of target genes; these speckle-associating target genes experience boosted induction upon p53 activation and belong to distinct functional categories compared to non-associating p53 target genes^21^. To determine if similar principles were true for HIF-2α-mediated DNA-speckle association, we performed RNA-seq at 3, 6, 9, and 12 hours after PT2399 treatment in 786-O cells (**Fig S2H**). We found that genes exhibiting decreased speckle association also showed a general decrease in gene expression at 3 hours, which became more pronounced at later time points (**Fig 2H**, 2007 genes, compare to log2 fold change of 0 red line). We then evaluated how many HIF-2α-responsive genes, as defined by reduced RNA expression at any time point upon PT2399 treatment, showed HIF-2α-regulated DNA-speckle association, as measured by SON TSA-seq or Cut&Run (**Fig 2I**). This analysis showed significant overlap between HIF-2α target genes and genes with decreased SON signal upon PT2399 treatment, revealing that ∼25% of HIF-2α target genes experience PT2399-mediated decreases in speckle association. Thus, like p53, HIF-2α regulates speckle association of a subset of its target genes.

To evaluate whether speckle-associating and non-speckle-associating HIF-2α target genes differ in their level of expression, we examined RNA-seq data of these two gene groups, with 250 HIF-2α target genes that had decreased SON in TSA-seq or Cut&Run with PT2399 (padj < 0.01) defined as “associating”, and 522 HIF-2α target genes that did not have decreased SON in TSA-seq or Cut&Run (padj > 0.1) defined as “non-associating”. We found that speckle-associating HIF-2α target genes tended to have higher RNA expression at baseline without HIF-2α inhibition (**Fig 2J**, left; DMSO treated 786-O cells), and showed slower reduction in gene expression upon PT2399 treatment compared to non-speckle-associating targets (**Fig 2J**, right). Hence, HIF-2α target genes that experience HIF-2α-mediated speckle association are more highly expressed and show distinct responses to PT2399 HIF-2α inhibition.

We next assessed whether speckle-associating versus non-associating HIF-2α target genes belonged to distinct functional categories using Gene Ontology (GO) analysis. We found that speckle-associating HIF-2α target genes were specifically enriched in terms related to inflammation (e.g. IkB/NFkB signaling and cell response to IL4), protein transport, metabolism, and cell division (**Fig 2K**, left), while non-speckle-associating HIF-2α target genes were specifically enriched in terms related to cell differentiation and tissue development, importantly including many terms relating to angiogenesis (**Fig 2K**, right). These findings indicate that nuclear speckles act upon a specific subset of HIF-2α target genes.

### The HIF-2α speckle targeting motifs (STMs) are essential for HIF-2α-driven DNA-speckle association and HIF-2α induction of speckle-associating genes

Our above experiments utilize PT2399 to globally lower HIF-2α function. We next sought to specifically disrupt speckle targeting abilities of HIF-2α by deleting its two putative speckle targeting motifs (STMs; see **Fig 2A**). We employed dox-inducible HIF-2α with or without intact STMs (HIF-2α ref-seq NP_001421.2 Δ455-470; Δ776-791, called ΔSTMs) in MCF7 cells, a breast cancer cell line with low endogenous HIF-2α protein levels (**Fig 3A**; **Fig S3A**, left). To prevent pVHL-mediated proteosomal degradation of ectopic HIF-2α, we included two proline substitution mutations (P405A and P531A, termed HIF2As for stabilized HIF-2α, which do not overlap with STMs)^32,36–38^. We selected three single-cell clones of varying HIF-2α amounts in HIF2As-wtSTM and HIF2As-ΔSTM (**Fig 3B**; **Fig S3A** *selected clones), and found via immunoDNA-FISH that dox-induction of HIF2As-wtSTM led to closer speckle association of HIF-2α target *DDIT4*, and, in contrast, no change in control neighbor gene *DDX21* (**Fig 3C**; compare orange HIF2As-wtSTM with grey no HIF-2α negative control). As the key comparison, across the same range of HIF-2α levels, HIF2As-ΔSTMs failed to induce speckle association of *DDIT4* (**Fig 3C**; compare blue HIF2As-ΔSTMs with orange HIF2As-wtSTM and grey no HIF-2α control). These results demonstrate that the STMs are necessary for HIF-2α-mediated target gene-speckle association.

**Figure 3.**
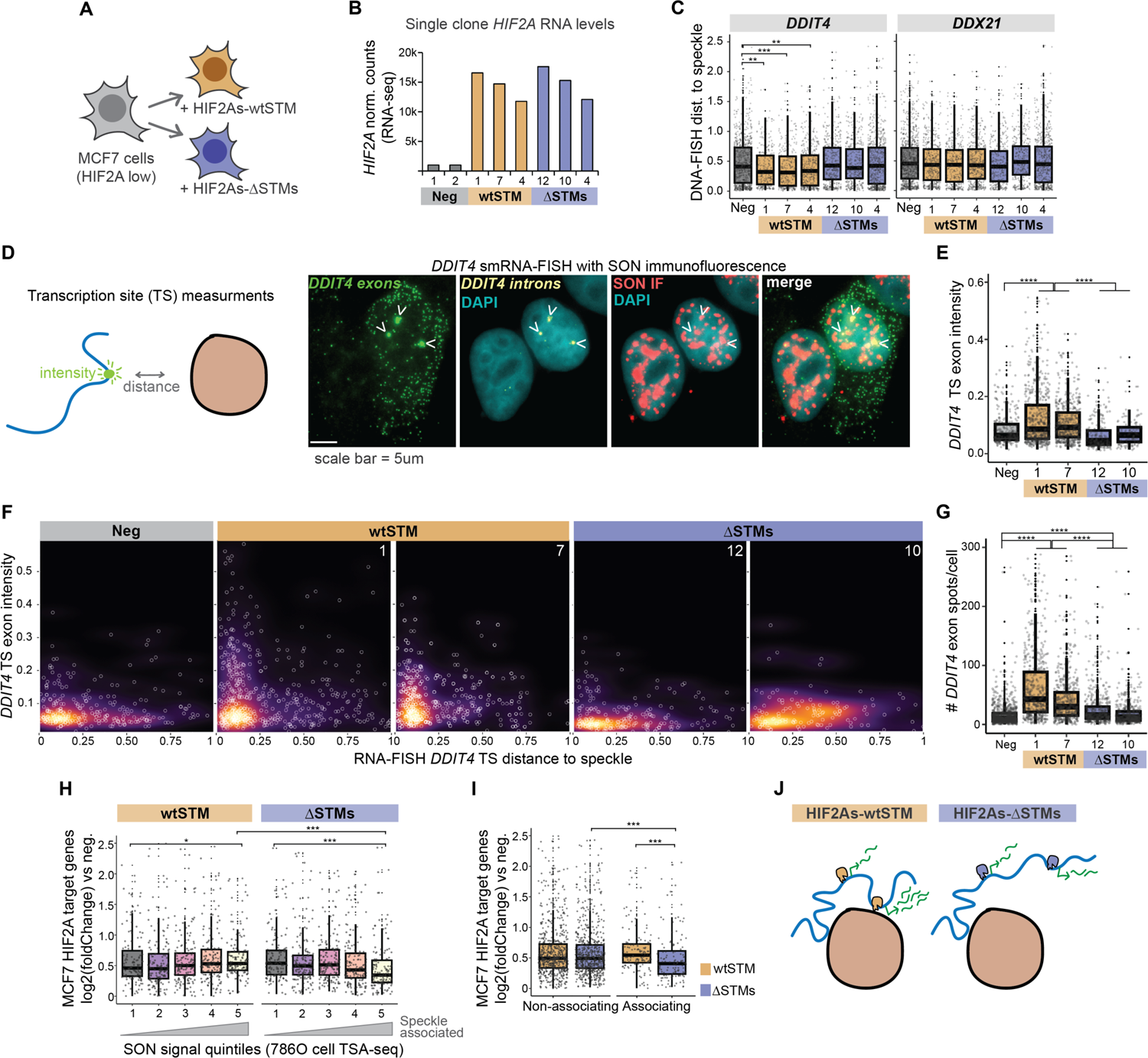
The HIF-2α speckle targeting motifs (STMs) are essential for HIF-2α-driven DNA-speckle association. A) Schematic of experiment. HIF2As – stabilized HIF-2α with P405A;P531A. wtSTM – speckle targeting motifs intact. ΔSTMs – speckle targeting motifs deleted: Δ455-470; Δ776-791. B) RNA-seq normalized counts *HIF-2α* in negative control (grey) and single-cell clones of dox-induced HIF2As-wtSTM (yellow) and HIF2As-ΔSTMs (blue). See also **Fig S3A**. C) immunoDNA-FISH loci distance to speckle for *DDIT4* and *DDIT4* in negative control and three single-cell clones of dox-induced HIF2As-wtSTM (yellow) or HIF2As-ΔSTMs (blue). D) Schematic of immunoRNA-FISH and example image of *DDIT4* exons (green), *DDIT4* introns (yellow), DAPI (cyan), and SON (red). Arrowheads indicate location of *DDIT4* transcription sites. E) *DDIT4* transcription site (TS) exon intensities from immunoRNA-FISH data. Each dot is an individual transcription site. See **Fig S3C** for intron intensities. F) Relationship between *DDIT4* transcription site (TS) exon intensities and distance to speckle. Each point is an individual transcription site. See **Fig S3B** for TS intron intensity plots. G) Number of *DDIT4* exon spots per cell. Each dot is an individual cell. H) Fold change of MCF7 HIF-2α target genes with dox-induction of HIF2As-wtSTM or HIF2As-ΔSTMs split into quintiles by speckle association status from Fig 2. Each dot is one HIF-2α target gene. I) Fold change as in **H** split by whether the target gene did (“Associating”) or did not (“Non-associating”) decrease speckle association in 786-O cells treated with PT2399 from Fig 2. Each dot is one HIF-2α target gene. J) Model showing loss of speckle association and expression with HIF-2α STM deletion.

The speckle-targeting-defective HIF2As-ΔSTMs mutant provides a specific tool to evaluate the gene expression consequences of disrupting DNA-speckle association. We employed single molecule RNA-FISH (smRNA-FISH) of *DDIT4* in the above MCF7 cells (with doxycycline-induced HIF2As-wtSTM or HIF2As-ΔSTM) using intron and exon probe sets which enable transcription site exon probe intensity to be measured concurrently with transcription site distance to the nearest speckle (**Fig 3D**, diagram on left). Transcription site intensities increased upon application of HIF2As-wtSTM, whereas they failed to increase in HIF2As-ΔSTMs (**Fig 3E**). We compared transcription site exon intensity to speckle distances, and observed a characteristic L-shaped distribution (also observed in our p53 results in ^21^) with RNA building up at speckle-adjacent sites in cells with HIF2As-wtSTM, but not in cells without induction of HIF-2α or with HIF2As-ΔSTMs (**Fig 3F**). We observed the same trends with *DDIT4* intronic probes (**Fig S3B-C**), suggesting that HIF-2α STMs are essential for production of nascent unspliced *DDIT4* RNAs within transcription sites. We quantified the number of exonic spots to estimate total *DDIT4* RNA molecules per cell, revealing that HIF2As-ΔSTMs greatly lowered induction of *DDIT4* compared to HIF2As-wtSTM (**Fig 3G**). These results indicate that intact speckle-targeting of HIF-2α is required for expression of speckle-associating HIF-2α target gene, *DDIT4*.

We then evaluated the genome-wide consequences of HIF-2α STM deletions via RNA-seq, testing whether disrupting speckle targeting had specific consequences based on speckle association. We found that HIF2As-ΔSTMs was specifically defective at inducing speckle-associated HIF-2α target genes (**Fig 3H**; yellow bin 5 – most speckle associated genes), while non-speckle-associated HIF-2α target genes were similarly induced by HIF2As-wtSTM and HIF2As-ΔSTMs (**Fig 3H**; grey bin 1 – least speckle associated genes).

Similarly, by comparing induction of HIF-2α target genes that did or did not have speckle association regulated by PT2399 (from **Fig 2**), we found that HIF2As-ΔSTMs was defective at inducing genes that had speckle association regulated by HIF-2α (**Fig 3I**; ΔSTMs blue “non-associating” versus ΔSTMs blue “associating”), whereas HIF2As-wtSTM showed comparable induction of HIF-2α target genes that did or did not have HIF-2α-dependent changes in speckle association (**Fig 3I**; wtSTM orange “Non-associating” versus wtSTM orange “Associating”). These results demonstrate that deletion of HIF-2α STMs shifts the balance of HIF-2α target gene induction, with specific consequences for speckle-associated target genes.

In aggregate, we identified a motif for speckle association (the STM) that reoccurs among many key regulators of gene expression, such as HIF-2α. Our results show (1) that HIF-2α works via its STMs to accomplish DNA-speckle association, and (2) that disruption of HIF-2α speckle targeting ability compromises induction of the speckle-associating subset of its target genes. These findings together support a model whereby nuclear speckles and DNA-speckle targeting represent a distinct layer of gene regulation that shifts the balance of genes that are preferentially expressed. We hypothesize that this layer of gene regulation can be manipulated by altering the transcription factor speckle targeting abilities, as with HIF-2α STM deletions above, or by alterations in the speckles themselves, as with the cancer speckle signature (see **Fig 1**), explored further below.

### Speckle Signature I tumors are enriched in oxidative phosphorylation and ribosome pathways, with higher RNA expression of speckle-associating megabase-sized gene neighborhoods

The speckle signature, while present in many cancers, was particularly predictive of survival in ccRCC (see **Fig 1**). Given our findings that HIF-2α regulates DNA-speckle association, we hypothesized that HIF-2α combines with speckle phenotypes, resulting in poor ccRCC outcomes. To first broadly understand the consequences of cancer speckle signature, we shifted our attention to deeper analysis of gene expression differences between speckle signature patient groups in TCGA data. We divided TCGA samples into Signature I and Signature II groups using the top and bottom 25% of sample speckle scores (from **Fig 1**; Signature I – top 25%, Signature II – bottom 25%), calculated gene expression fold changes, and used Gene Set Enrichment Analysis (GSEA) to identify which biological pathways were altered between the two patient groups. We found striking enrichment of “Oxidative phosphorylation” and “Ribosome” in the Signature I group among all cancer types, including ccRCC (KIRC) (**Fig S4A** – example GSEA plots; **Fig 4A** – Hallmark pathways; **Fig S4B** – KEGG pathways). Hence, across many cancer types, speckle Signature I correlates with increased oxidative phosphorylation and ribosomal pathways, supporting that speckle Signature I tumors, which reflect the aberrant speckle signature acquired in cancer, may exist in a “hyper-productive” state with enhanced metabolic and protein production capacity.

**Figure 4.**
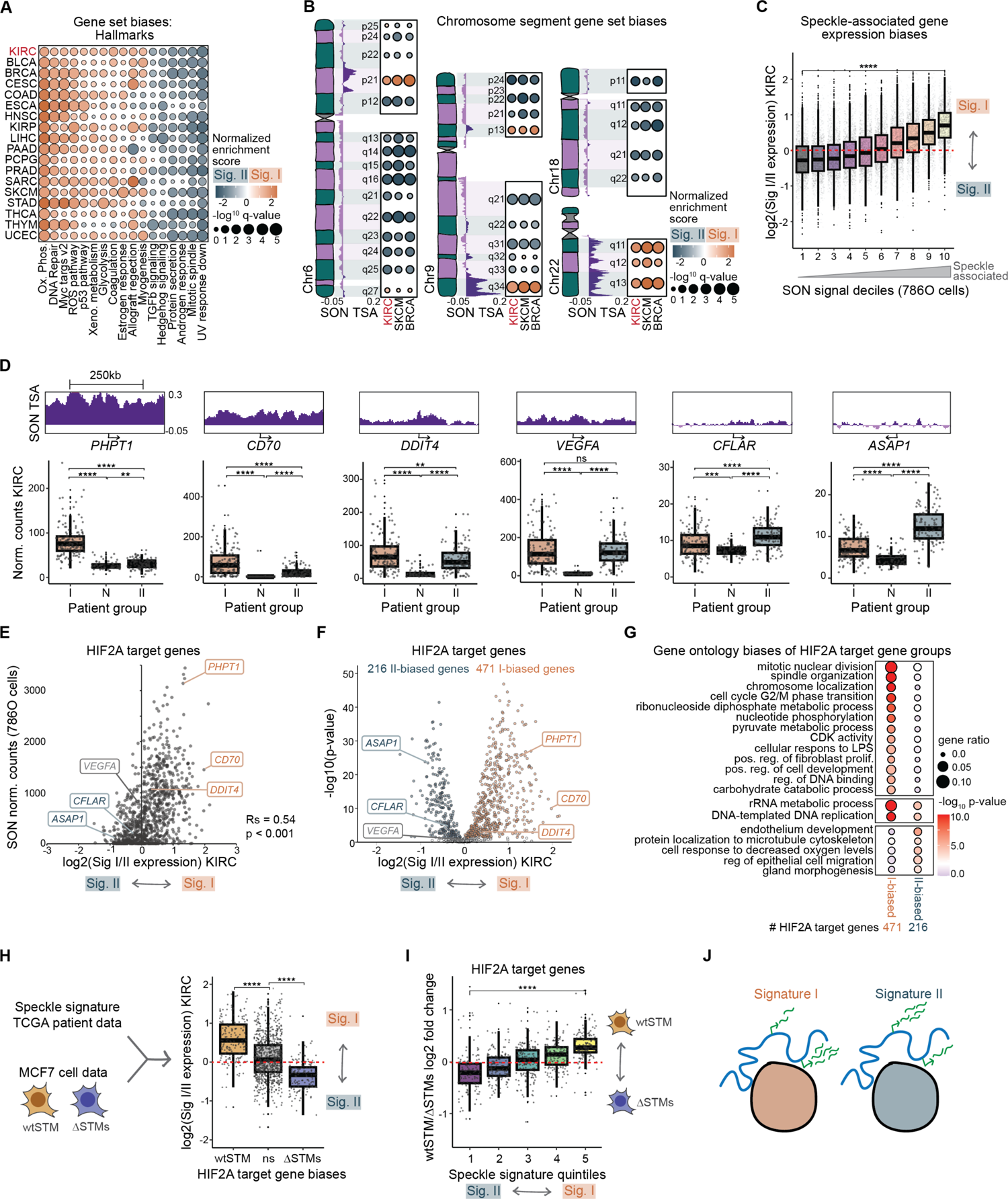
Cell line-mapped speckle associated genes are more highly expressed in the speckle Signature I patient group. A) Hallmark gene set enrichment statistics for Signature I versus Signature II speckle patient groups. ccRCC (KIRC) is in red text. See also **Fig S4A-B**. B) Depictions of chromosomes, SON TSA-seq genome browser tracks, and cytogenetic band enrichment statistics of Signature I versus Signature II for chromosomes 6, 9, 18, and 22. See also **Fig S4A**, right and **Fig S4C**. C) Decile plot split by SON TSA-seq signal showing Signature I to Signature II expression ratio in the KIRC TCGA cohort. Each dot represents one gene. Significance calculated using Wilcoxon test. D) SON TSA-seq genome browser tracks of HIF-2α target genes with varying speckle association (top, purple) and expression in Signature I (I), normal adjacent tissue (N), and Signature II (II) samples from the TCGA KIRC cohort. Each dot represents one sample. Significance calculated using Wilcoxon test. E) SON normalized counts versus speckle patient group gene expression for HIF-2α target genes. Rs – R Spearman’s correlation coefficient. p-value represents correlation significance. F) Volcano plot of speckle patient group gene expression ratio for HIF-2α target genes versus -log10(p-value) statistic for gene expression difference. G) Gene ontology showing HIF-2α pathways biased toward Signature I (top) or Signature II (bottom) speckle patient group. H) Schematic depicting combination of TCGA speckle signature data with STM deletion data in MCF7 cells from Fig 3 (left). Boxplot of HIF-2α target genes showing Signature I to Signature II expression ratio in the KIRC TCGA cohort split by expression differences between HIF2As-wtSTM and HIF2As-ΔSTMs. I) Quintile plot of HIF-2α target gene fold changes of HIF2As-wtSTM versus HIF2As-ΔSTMs split by speckle signature bias. J) Model depicting lower expression of speckle-associated genes in speckle Signature II. ** – p < 0.01; *** – p < 0.001; **** – p < 0.0001, ns – not significant SON TSA-seq data in **B**, **C**, **D** and **E** is from Fig 2 786-O DMSO condition.

We also performed positional gene set analysis, which consists of gene sets corresponding to human chromosome cytogenetic bands, delineating megabase-sized neighborhoods of heterochromatin or euchromatin on the linear genome^39^. Our analysis revealed astonishing expression biases of certain chromosome cytogenetic bands. For example, genes in the CHR1P31 band were highly enriched in the speckle Signature II patient group, while genes in CHR19Q13 were highly enriched in the speckle Signature I patient group (**Fig S4A**, right-most plots). We plotted cytogenetic band gene set enrichment statistics on entire chromosomes side-by-side with SON TSA-seq (from **Fig 2**), finding that Signature I-biased bands corresponded to regions of the genome with higher SON signal (**Fig 4B** and **Fig S4C**; e.g. CHR6P21, CHR9Q34, all of Chr22). Reciprocally, Signature II-biased bands corresponded to regions of the genome with lower SON signal (**Fig 4B** and **Fig S4C**; e.g. Q arm of Chr6, all of Chr18). These data suggest that speckle signature affects large groups of genes depending on speckle association.

We quantified the per-gene relationship between speckle association and expression in the speckle signature patient groups. We separated all expressed genes into deciles depending on their speckle association and graphed the expression fold change of the speckle Signature I versus II patient groups for each gene (**Fig 4C**). In agreement with our cytogenetic band enrichment results, this analysis revealed that speckle associated genes were biased towards higher expression in the Signature I patient group (**Fig 4C**; yellow bin 10, compare to log2 fold change of 0, red line), and the genes that were most speckle-distal were biased towards higher expression in the Signature II patient group (**Fig 4C**; grey bin 1). These data reveal a propensity of speckle associated genes to be more highly expressed in speckle Signature I samples, suggesting that Signature I speckle components are positive regulators of gene-activating speckle functions (explored further in **Fig 5**).

**Figure 5.**
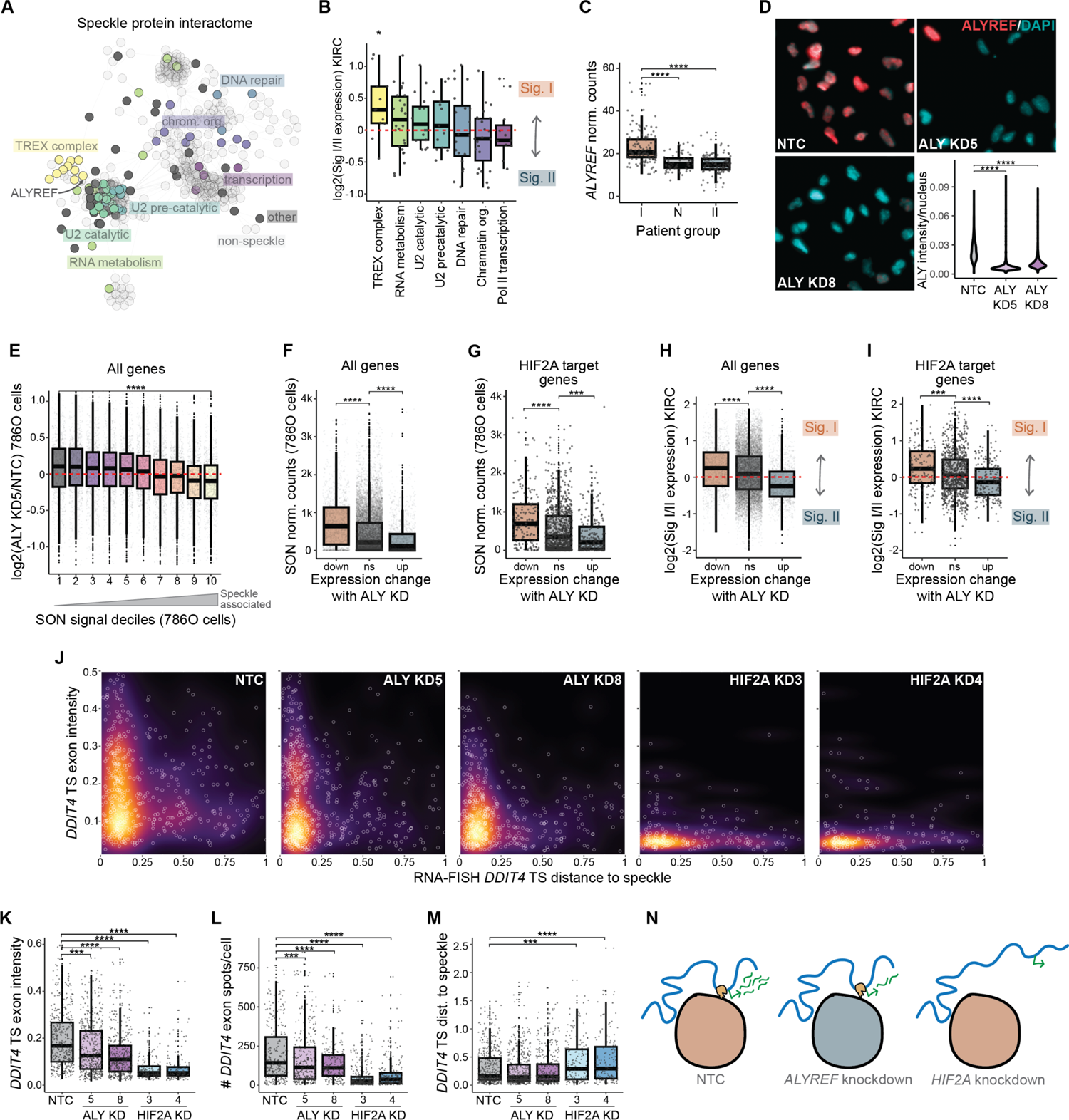
TREX complex member, ALYREF, is required for full expression of speckle associated genes. A) Speckle protein physical interactome network. Transparent nodes are non-speckle interactors. Opaque nodes are speckle-resident proteins and are colored based on functional annotation. B) Signature I to Signature II expression for speckle protein genes within the designated functional categories. Significance calculated using T-test versus null hypothesis of 0. C) Expression of *ALYREF* in Signature I (I), normal adjacent tissue (N), and Signature II (II) sample groups from the TCGA KIRC cohort. D) ALYREF immunofluorescence (red) and DAPI (cyan) in 786-O cells treated with non-targeting (NTC) or *ALYREF* siRNAs (ALY KD5 and ALY KD8), and quantification of ALYREF signal intensity (violin plot). E) Decile plot fold change upon *ALYREF* knockdown (ALY KD5; ALY KD8 in **Fig S5A**) split into speckle association deciles. F) SON TSA-seq signal of genes that decrease upon *ALYREF* knockdown (orange, “down”), are unchanged (grey, “ns”), and increase (blue, “up”).G) Same as **F** for HIF-2α target genes H) Signature I to Signature II expression ratio in the KIRC TCGA cohort for genes that decrease upon *ALYREF* knockdown (orange, “down”), are unchanged (grey, “ns”), and increase (blue, “up”). I) Same as **H** for HIF-2α target genes. J) Single-molecule immunoRNA-FISH quantification of *DDIT4* transcription site (TS) distance to speckle versus exon intensity for non-targeting control (NTC), *ALYREF* (ALY KD5 and ALY KD8), and *HIF-2α* (HIF-2α KD3 and HIF-2α KD4) siRNA treatment in 786-O cells (see **Fig S5B** for Western blot of *HIF-2α* knockdown). Each point represents an individual transcription site. K) *DDIT4* transcription site exon intensities with *ALYREF* and *HIF-2α* knockdown (same data as **J**). Each dot represents an individual transcription site. L) Number of *DDIT4* exon spots per cell. Each dot represents an individual cell. M) *DDIT4* transcription site distance to nearest speckle (same data as **J**). Each dot represents an individual transcription site. N) Model showing decreased expression of speckle associated genes upon *ALYREF* knockdown. Significance calculated using Wilcoxon tests. * – p < 0.05; ** – p < 0.01; *** – p < 0.001; **** – p < 0.0001, ns – not significant SON TSA-seq data in **E**, **F**, and **G** is from Fig 2 786-O DMSO condition.

### Signature I and Signature II-biased HIF-2α target genes reflect distinct functional categories

We next focused on HIF-2α target genes within the speckle signature analysis to elucidate consequences of speckle alterations in ccRCC. Compared to normal tissues, HIF-2α target genes were more highly expressed in both speckle Signature I and Signature II patient groups (**Fig 4D**; N – normal adjacent tissue samples; I – speckle Signature I samples; II – speckle Signature II samples). However, certain target genes were more highly expressed in the Signature I patient group (e.g. *PHPT1* in **Fig 4D**, left), while others were more highly expressed in the Signature II patient group (e.g. *ASAP1* in **Fig 4D**, right). These expression biases corresponded to differences in speckle association, shown at representative genes (**Fig 4D**, top SON TSA-seq tracks), and for all HIF-2α target genes (**Fig 4E**). In total, 471 of 986 HIF-2α target genes were more highly expressed in the Signature I patient group, and 216 were more highly expressed in the Signature II patient group (**Fig 4F**). Using Gene Ontology analysis, we found that Signature I-biased HIF-2α target genes were enriched in cell cycle, metabolism, and inflammatory pathways, and Signature II-biased HIF-2α target genes were enriched in endothelium development (angiogenesis) (**Fig 4G**). Importantly, these functional terms were similar to the terms we found as differential between HIF-2α target genes that did or did not have HIF-2α regulated speckle association, with shared pathways between Signature I-biased genes and speckle-associating genes as well as between Signature II-biased genes and non-speckle-associating target genes. (compare **Fig 2K** with **Fig 4G**). These findings indicate that HIF-2α functional programs in ccRCC correlate with speckle signature. We propose that these speckle-relevant HIF-2α target gene expression biases may underly the speckle-based differences in patient outcomes in ccRCC.

### Loss of HIF-2α DNA-speckle targeting phenocopies HIF-2α target gene expression biases of the better-outcome speckle Signature II patient group

Our data thus far indicates that HIF-2α regulates expression of speckle-associating target genes by mediating their speckle association. Meanwhile, our correlational evidence supports that speckles regulate expression of speckle-associating HIF-2α target genes by changing the gene activation functions of nuclear speckles. We hypothesized that HIF-2α target genes differentially expressed between the two speckle signatures were specifically and similarly dependent on the HIF-2α speckle targeting motif, STM. We thus evaluated HIF-2α target gene expression biases between speckle signatures in the context of HIF-2α target genes that were differentially regulated between HIF2As-wtSTM and HIF2As-ΔSTMs. Indeed, nearly all the HIF-2α target genes that were preferentially expressed in HIF2As-wtSTM were more highly expressed in the speckle Signature I patient group (**Fig 4H**; orange box, above 0 means higher in Signature I patient group). In contrast, the HIF-2α target genes that were preferentially expressed in HIF2As-ΔSTMs were more highly expressed in the speckle Signature II patient group (**Fig 4H**; blue box, below 0 means higher in Signature II patient group); meanwhile, HIF-2α target genes that did not have a wtSTM/ΔSTMs preference showed a mixture of I and II-biased expression patterns (**Fig 4H**, grey). We also separated HIF-2α target genes into quintiles based on speckle Signature I or II gene expression biases and graphed fold change of HIF2As-wtSTM versus HIF2As-ΔSTMs (**Fig 4I**). This analysis demonstrated that the most Signature I-biased HIF-2α target genes had the highest expression in the HIF2As-wtSTM condition (**Fig 4I**; yellow bin 5, above 0 means higher in HIF2As-wtSTM), and the most Signature II-biased HIF-2α target genes had the highest expression in the HIF2As-ΔSTMs condition (**Fig 4I**; purple bin 1; below 0 means higher in HIF2As-ΔSTM). Hence, there was a strong concordance between gene expression differences between speckle signature patient groups and changes observed upon disrupting HIF-2α speckle targeting abilities in tissue culture. Speckle Signature I corresponds to boosted expression of speckle-associated HIF-2α targets and mimics the effects of HIF-2α that is speckle-targeting-competent (wtSTM), whereas speckle Signature II corresponds to boosted expression of non-speckle-associating HIF-2α targets and mimics the effects of HIF-2α that is speckle-targeting-defective (ΔSTMs) (see models in **Fig 3J** and **Fig 4J**). Thus, our results show striking parallels of HIF-2α target gene expression upon differential speckles in human tumors and differential HIF-2α DNA-speckle targeting in tissue culture, supporting a model whereby HIF-2α target gene expression can be similarly regulated by changes in speckles or DNA-speckle association.

### The TREX complex is elevated in speckle Signature I patients and TREX complex member ALYREF is required for expression of speckle-associated genes

Our results link speckle Signature I to higher expression of speckle associated genes (**Fig 4B-C**), leading us to investigate whether variation in the speckle composition of Signature I compared to Signature II might explain gene expression differences. To reveal the protein complexes and functional categories of proteins that reside within speckles and predominate within the speckle signatures, we utilized speckle resident protein annotations from the Human Protein Atlas and physical interactions from STRING-DB to perform network analysis of speckle resident proteins and their interactors. This analysis revealed hubs of interconnected proteins that represented distinct protein complexes, providing a comprehensive view of complexes residing within speckles (**Fig 5A**; speckle-resident proteins are solid colors, non-speckle-resident interactors are transparent, only connected nodes are displayed). Using STRING-DB to identify the key functional annotations, we classified seven major categories of speckle-resident proteins: RNA metabolism, DNA repair, transcription, chromatin organization, splicing (subdivided into U2 pre-catalytic and U2 catalytic complexes), and the transcriptional export (TREX) complex (**Fig 5A**). Examining RNA expression differences between Signature I and Signature II speckle patient groups, we found that 8 of 10 speckle-resident TREX complex members were more highly expressed in the Signature I group, while the other functional groups were approximately evenly split between the two signatures (**Fig 5B**; yellow box). These results highlight TREX as a potentially key differential protein complex that is more highly expressed in the speckle Signature I patient group.

Elevated TREX in speckle Signature I patients led us to hypothesize that the TREX complex may contribute to the gene-activating functions of nuclear speckles. TREX plays a central role in gene expression, accompanying mRNAs from transcription to nuclear export (reviewed in ^40^). Within TREX, the ALYREF protein is a central component that is loaded onto mRNAs co-transcriptionally, promoting pre-mRNA processing and export, and we find that it was elevated in Signature I tumor samples (**Fig 5C**). Depletion of ALYREF alters gene expression and reduces Poll II gene occupancy^41^, supporting an additional role for ALYREF in transcription efficiency. To investigate a correlation between requirement of ALYREF for speckle-associated expression, we used siRNA to knockdown *ALYREF* in 786-O ccRCC cells (**Fig 5D**). We found that the most speckle associated genes (bin 10, yellow) were the most decreased in expression upon *ALYREF* knockdown, while the least associated genes were the most increased in expression (bin 1, grey) (**Fig 5E**, *ALYREF* siRNA #5; **Fig S5A**, *ALYREF* siRNA #8). We also found higher levels of speckle association of genes significantly decreasing in expression upon *ALYREF* knockdown compared to unchanging genes and significantly higher genes (**Fig 5F**). This trend existed genome-wide (**Fig 5F**), and was true of HIF-2α target genes (**Fig 5G**). These results indicate that ALYREF is required for expression of speckle-associated genes.

Because *ALYREF* is more highly expressed in the speckle Signature I patient group (**Fig 5C**), we hypothesized that knockdown of *ALYREF* may exhibit similar gene expression biases to the speckle Signature. Consistent with this, we found that genes decreasing with *ALYREF* knockdown were more highly expressed in the speckle Signature I patient group, while genes increasing with *ALYREF* knockdown were more highly expressed in the speckle Signature II patient group (**Fig 5H**). This was also observed for HIF-2α-responsive genes (**Fig 5I**). Hence, *ALYREF* knockdown in 786-O ccRCC cells phenocopies gene expression differences of ccRCC speckle signature patient groups.

To further evaluate the relationship between ALYREF and speckle-associated gene expression, we performed smRNA-FISH using *DDIT4* HIF-2α target gene probes (as in **Fig 3**). As a reference point for disrupting *DDIT4* expression and speckle association, we compared *ALYREF* knockdown to HIF-2α knockdown in 786-O cells (**Fig S5B** shows confirmation of HIF-2α knockdown). We found that *ALYREF* knockdown partially compromised RNA levels within *DDIT4* transcription sites (**Fig 5J**, compare NTC and ALYkd panels; quantified in **Fig 5K**), with lower total *DDIT4* RNA levels (**Fig 5L**). *ALYREF* knockdown did not affect *DDIT4* speckle association (**Fig 5M**). Hence, *ALYREF* knockdown compromised expression of speckle associated genes without influencing speckle association status (**Fig 5N**). These results suggest that ALYREF contributes to the ability of nuclear speckles to boost gene expression but is not required for HIF-2α-mediated association of genes with speckles.

### Nuclear positioning of the SON speckle marker predicts patient outcomes in ccRCC

Our findings with *ALYREF* knockdown recapitulated some of the speckle signature group differences in ccRCC tumor samples. We also noted changes in nuclear speckle morphology, including fewer, but larger, speckles with *ALYREF* knockdown and a slight reduction in the amount of SON signal at the nuclear periphery (**Fig S5C-G**). These findings prompted us to evaluate whether ccRCC tumor samples show variation of imaging-based speckle phenotypes. To this end, we obtained a tumor microarray that contained 86 ccRCC formalin-fixed paraffin-embedded (FFPE) tumor sections with matched adjacent tissue and associated patient survival data. We performed SON immunofluorescence and scanned the entirety of each tissue section at 20x magnification (**Fig 6A**, left). Using CellProfiler, we identified nuclei based on DAPI staining and calculated per-nucleus measurements of SON texture, intensity, and radial distribution. For radial distribution, the nucleus was divided into four concentric bins and the proportion of immunofluorescence signal residing in each bin was calculated (**Fig 6B**, right schematic). Based on the median sample value of each measurement, we separated ccRCC tumors into the top and bottom half and performed Kaplan-Meier analysis for each measurement (**Fig 6A**, right schematic). Plotting Kaplan-Meier statistical measures, we found that SON radial distribution correlated most significantly with survival (**Fig 6B**, yellow points). Specifically, the most significant predictors of survival were fraction of SON in the most central nucleus bin followed by fraction of SON in the most peripheral nucleus bin (**Fig 6B**, top two labelled yellow points). These differences were clearly apparent in tumor images: high central SON tumors showed speckles in the center of nuclei (**Fig 6C**; C11 and I15 tumors), and high peripheral SON tumors showed speckles spread throughout the nuclei (**Fig 6C**; C9 and E17 tumors). As these are reciprocal measurements, samples with low central and high peripheral SON reflect the same patient group. Tumors with low central (high peripheral) SON were associated with worse patient outcomes compared to tumors with high central (low peripheral) SON (**Fig 6D-E**). These results reveal that imaging measurements of speckle phenotypes predict outcomes in ccRCC.

**Figure 6.**
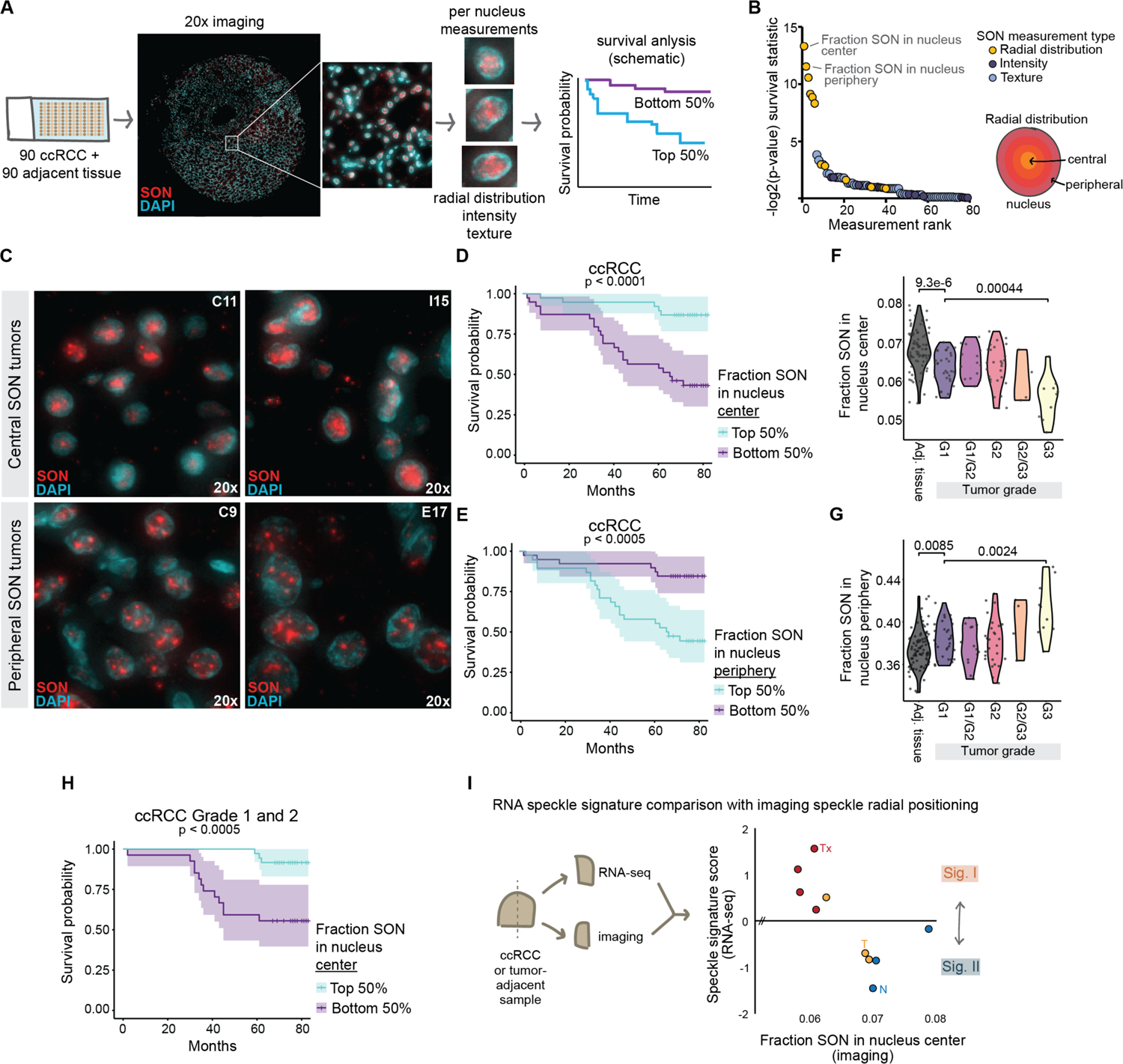
Imaging-based SON radial positioning with the nucleus predicts ccRCC outcomes and strongly correlates with RNA-based speckle signature. A) Schematic of ccRCC tumor array (left), SON immunofluorescence (red) and DAPI (cyan) imaging, image analysis, and survival calculation (right). B) Left; -log2 Kaplan Meier statistic for each SON imaging measurement, colored by type of measurement. Right; schematic of radial distribution measurement. C) Representative 20x images of tumors with high central SON (top) and high peripheral SON (bottom). D) Kaplan Meier plot of ccRCC split by the top and bottom 50% of SON in the nucleus center. E) Same as **D** for SON in the nucleus periphery. F) Fraction of SON in the nucleus center for adjacent tissue (grey) and ccRCC tumors split by tumor grade. Each dot represents the median value of all the nuclei measured in one sample. Significance calculated by Wilcoxon test. G) Same as **F** for SON in the nucleus periphery. H) Kaplan Meier plot of Grade 1 and 2 ccRCC split by the top and bottom 50% of SON in the nucleus center. I) Relationship between speckle signature score and the fraction of SON in the nucleus center from ccRCC tumor and adjacent normal samples in split for RNA and imaging (as in schematic, left). Tx – Xenograft tumor from mice; all four are from same individual donor, different mice. T – primary tumor. N – tumor-adjacent normal samples.

We compared ccRCC samples to matched normal adjacent tissues and separated ccRCC based on tumor grade. This showed that normal adjacent tissues have higher central SON (lower peripheral SON) compared to ccRCC, and SON becomes less central (more peripheral) in later grade ccRCC (**Fig 6F-G**). We evaluated whether altered radial distribution of SON in higher grade tumors strongly influenced survival correlations, performing Kaplan-Meier analysis on the subset of Grade1 and 2 tumors, which had similar SON nuclear distributions. Despite excluding higher grade ccRCC, radial positioning of SON was predictive of ccRCC outcomes (**Fig 6H**), indicating that nuclear speckle phenotypes predict outcomes even in early-stage ccRCC, providing potential diagnostic relevance.

We next directly assessed both RNA- and imaging-based measurements of speckle phenotype in the same cohort of samples, hypothesizing that samples with lower central SON would correspond to Signature I speckle protein gene expression (see **Fig 1**). We obtained clinical ccRCC tumor and tumor-adjacent samples and divided them for RNA-seq and FFPE SON immunofluorescence, including three tumor-adjacent primary tubule renal epithelium samples, three primary human ccRCC tumors, and four mouse xenograft-derived ccRCC tumors derived from the same individual. We calculated speckle protein gene expression scores via RNA-seq as previously (**Fig 1**). As predicted, primary renal tubule epithelial samples (normal adjacent) displayed Signature II RNA speckle scores and high central SON by imaging (**Fig 6I**; blue points; N – normal primary renal tubule epithelium). Two primary ccRCC tumors also showed Signature II speckle scores with high central SON (**Fig 6I**; yellow points; T – primary tumor). In contrast, the remaining primary ccRCC tumor and all four patient-derived xenograft samples showed the opposite Signature I speckle scores, with corresponding low central SON, as predicted (**Fig 6I**; see yellow primary tumor point and red xenograft points (Tx) on upper left portion of graph). These direct comparisons strongly indicate that speckle Signature I manifests as more spread out and less central speckles associated with poor ccRCC survival (e.g. **Fig 6C**, bottom), while speckle Signature II manifests as more central larger speckles associated with better ccRCC survival (e.g., **Fig 6C**, top). Therefore, these data link RNA-seq and imaging-based speckle phenotypes, demonstrating that they may be used interchangeably to predict survival in ccRCC and suggesting that RNA-seq estimations of speckle phenotypes reflect *bona fide* differences in speckle morphologies, adding to potential therapeutic relevance.

## DISCUSSION

This study spotlights nuclear speckles as a distinct layer of gene regulation that skews HIF-2α functional programs and is correlated with poor patient outcomes in ccRCC. Our data indicate DNA-speckle targeting by transcription factors is a generalizable mechanism employed by transcription factors p53 and HIF-2α, via their speckle targeting motifs (STMs) defined herein. Identification of putative STMs that reoccur among gene regulatory proteins (**Fig 2**) suggests that speckle targeting may be a broadly utilized gene-regulatory mechanism. Based on mutagenesis of HIF-2α STMs (**Fig 3**), analysis of speckle variation in human samples (**Fig 4**), and manipulation of speckles in tissue culture (**Fig 5**), we reveal that nuclear speckles function by shifting the types of target genes that are preferentially induced by a transcription factor. We propose that this alteration of HIF-2α functional programs by speckles in ccRCC underlies speckle-based differences in patient survival.

While 23 other cancer types displayed similar speckle variation to ccRCC, none showed such dramatic survival differences (**Fig 1**). We speculate that the reason other cancer types do not show speckle-based survival differences may be due to more heterologous etiologies of other cancer types compared to ccRCC, which involves hyperactivation of HIF-2α in nearly all clinical cases^24^. Based on our findings that nuclear speckles skew HIF-2α functional programs, we hypothesize that speckle signature may be consequential for cancer subsets depending on specific transcription factor or gene expression dependencies. However, further studies are required to elucidate which specific cancer contexts may combine with nuclear speckle phenotypes to predict patient outcomes. Across most cancer types, the tumor adjacent normal tissue showed a speckle Signature II gene expression pattern (**Fig S1E**), suggesting that speckle Signature I represents an aberrant speckle state that is acquired in some tumors. The speckle Signature I patient group showed remarkable gene expression enrichment of ribosome and oxidative phosphorylation pathways in each cancer type (**Fig 4A** and **Fig S4B**), suggesting that speckle Signature I could reflect a hyper-productive cell state with enhanced metabolism and protein production capabilities. Given that some existing cancer therapies target protein synthesis pathways (e.g. mTOR inhibitors^42^), we propose that future studies should investigate whether speckle signature predicts patient responses to particular therapeutic agents, hypothesizing that, across many cancer types, the strong enrichment of ribosome protein gene expression in speckle Signature I tumors renders them more sensitive to mTOR inhibition or other therapies targeting metabolism and protein production.

Each cancer type also showed remarkable biases in chromosome position-based gene set enrichment, where entire groups of genes within a cytogenetically-defined chromosomal band were heavily skewed towards higher expression in the speckle Signature I patient group at speckle-associated chromosome bands, while non-speckle-associated chromosome bands were skewed towards higher expression in the speckle Signature II patient group (**Fig 4B** and **Fig S4A,C**). These findings provide compelling evidence for the long-standing model that nuclear speckles represent gene regulatory neighborhoods in higher eukaryotes^43–45^.

Together, the existence of the speckle signature in many cancer types, its strong correlation with certain functional pathways, and the extreme cytogenetic chromosome segment biases observed in our study all support the model that the speckle signature is a widespread phenomenon that coordinates functional pathways by influencing entire gene neighborhoods based on speckle association status.

We propose that nuclear speckles exist in two states (model in **Fig 7**). In one state (**Fig 7**, left panel), defined by Signature I speckle protein gene expression patterns, speckles are distributed throughout the nucleus (**Fig 6**), members of the TREX complex are more highly expressed (**Fig 5**), and expression of speckle-associated genes is elevated (**Fig 4**) (see model in **Fig 7**). In the other state, defined by Signature II speckle protein gene expression patterns that are more similar to normal tissues, speckles are located more centrally within the nucleus (**Fig 6**), members of the TREX complex exhibit lower expression (**Fig 5**), and non-speckle-associated gene expression is elevated (**Fig 4**). Reduction of the TREX complex member, ALYREF, suppressed expression of speckle associated genes, and shifted cells towards a more Signature II-like gene expression phenotype (**Fig 5**). Despite the observed decrease in speckle-associated gene expression with *ALYREF* knockdown, speckle association of the *DDIT4* gene was not perturbed (**Fig 5**), indicating that this disruption of speckle associated gene expression was due to altered speckle functions rather than an inability of genes to be speckle localized. This is a critical finding because it indicates that speckle phenotypes and DNA-speckle association are separate gene regulatory strategies used by cells.

**Figure 7.**
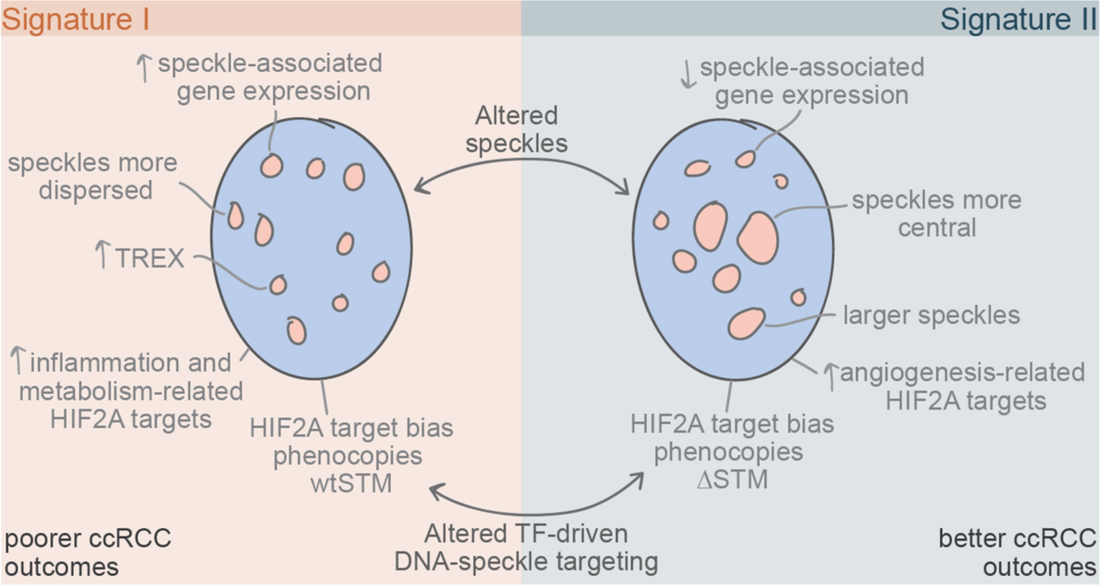
Model summarizing the two cancer speckle states. Our data supports that speckles exist in two states in cancer, with differential induction of speckle associated genes, speckle size and nuclear distribution, and speckle content. In ccRCC, speckle state correlates with patient survival and differential expression of HIF-2α functional pathways. Our findings indicate that speckle associated gene expression can be manipulated by changes in either speckles or transcription factor-driven DNA-speckle targeting.

While the speckle signature and *ALYREF* knockdown represent changes that broadly impact speckles, our experiments pinpointing the speckle targeting portions of HIF-2α provide a specific test for how speckles impact gene expression of associating DNA. Remarkably, the effects on HIF-2α target gene expression were paralleled by disruptions of HIF-2α speckle targeting abilities and differential ccRCC patient speckle signature (**Fig 4H-I**), supporting that disruption of DNA-speckle contacts or changes in speckles can shift the types of genes that are preferentially activated by a transcription factor. Speckle-associating and speckle Signature I-biased HIF-2α target genes were enriched in metabolism, inflammation, and cell cycle Gene Ontology categories, whereas non-speckle-associating and speckle Signature II-biased HIF-2α target genes were enriched in angiogenesis-related pathways (**Fig 2K** and **Fig 4G**). This functional shift of HIF-2α gene expression programs likely explain the striking survival differences based on speckle signature in ccRCC. However, further research is required to fully understand how these functional pathways link to different patient outcomes. Importantly, in ccRCC, several systemic therapies target HIF-2α and its downstream pathways^46^. Because our data suggests differential roles for HIF-2α depending on speckle signature, we speculate that speckles may be useful predictors for which therapies will be effective in particular ccRCC tumors.

Our identification of putative STMs within hundreds of gene regulatory proteins provide a platform for future investigations into the extent that DNA-speckle targeting is utilized by different factors. STMs occur in both direct DNA-binding transcription factors, as well as in many co-activators that engage with the genome by binding histones or DNA-binding proteins. Among these were several co-activators involved in enhancer functions, such as Histone K27 acetyltransferases, members of the Mediator complex, and Histone K4 methyltransferases. Given previous reports that enhancers and super-enhancers are highly enriched among speckle associated chromatin^14,15^, one hypothesis is that these STM-containing proteins work together to both mediate DNA-speckle association and promote deposition of activating histone modifications, creating a potent gene-activating environment where activating histone modifications are coordinated in nuclear space with the high concentration of RNA binding and processing factors found in nuclear speckles. Further research will be required in this area to disentangle enzymatic from speckle targeting functions of these co-activators, as well as to gain a full understanding of how histones, transcription, RNA processing, RNA stability, and RNA nuclear export may be coordinated at speckles to boost gene expression. At present, the mechanisms used by STMs to accomplish their functions are not known. In one model, STMs increase DNA affinity with nuclear speckles via interacting with a particular speckle-resident protein.

Alternatively, STMs may assist in trafficking DNA from other nuclear locations to speckles. Although a handful of speckle resident proteins contain STMs, we envision that speckle residency and speckle targeting are distinct molecular functions accomplished by different types of protein sequences. Other studies have identified protein components required for speckle residency, including a TP region of SF3B and the phosphorylated SR tracks of SR speckle resident proteins^47–49^. We note that these speckle residency protein domains do not overlap with the speckle targeting motifs identified in this study. While further efforts will be required to disentangle speckle targeting from speckle residency, we propose that STMs within speckle resident proteins may assist their speckle targeting, while other molecular features dictate speckle residency. Overall, our identification of STMs within both HIF-2α and p53 provides a molecular handle for future detailed inquiry of the molecular mechanisms underlying STMs and how they drive DNA-speckle association.

Overall, our findings illuminate nuclear speckles as underexplored regulators of gene expression that are consequential for human disease. We anticipate that our study of HIF-2α and ccRCC represents just one example, and that as nuclear speckles and DNA-speckle association by transcription factors become more broadly studied, additional examples of critical speckle functions in human health and disease will become evident.

## SUPPLEMENTAL FIGURE LEGENDS

**Figure S1.**
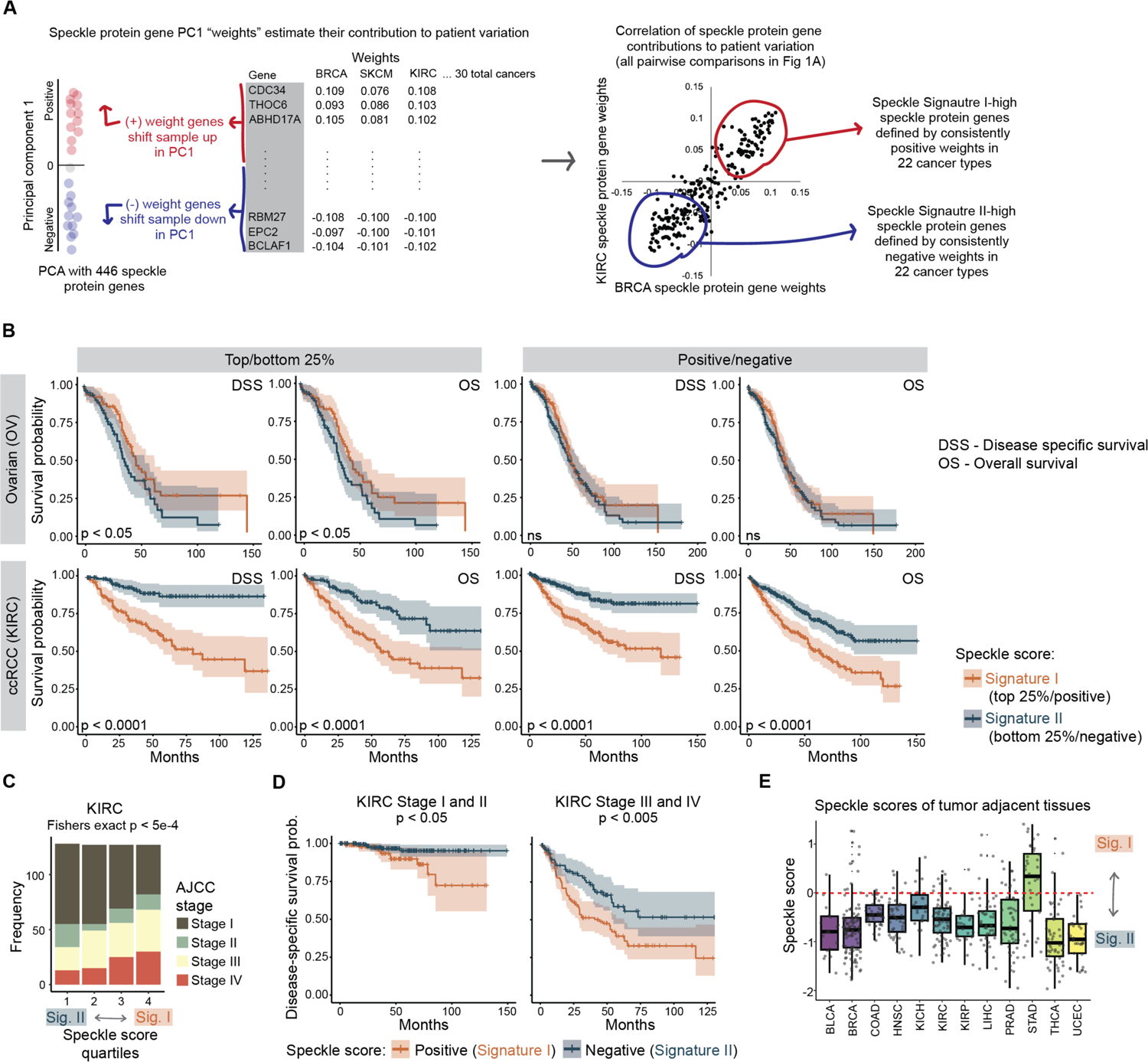
Supplement to Figure 1. A) Schematic showing how speckle protein genes with high contributions to patient variation were selected based on Principal Component 1 (PC1) weights. B) Kaplan Meier disease-specific (DSS) or overall survival (OS) analysis for ovarian cancer (OV, top) and ccRCC (KIRC, bottom) splitting individuals based on the top/bottom quartiles of speckle score (left) or on positive versus negative speckle score (right). C) Tumor AJCC stage frequency of ccRCC (TCGA KIRC cohort) tumors based on speckle scores split into quartiles. First quartile are the most Signature II tumors; fourth quartile are the most Signature I tumors. D) Kaplan Meier plots for early (left) and late (right) stage ccRCC (TCGA KIRC cohort). Patients were split into Signature I (positive speckle score) and Signature II (negative speckle score). E) Speckle scores of tumor adjacent normal tissues. Speckle scores are calculated independently for each tissue/cancer type. See **Table S1** for survival statistics for each cancer and Github (https://github.com/katealexander/speckleSignature.git) for detailed instructions, scripts used, and additional files generated relating to speckle signature calculations and survival analysis.

**Figure S2.**
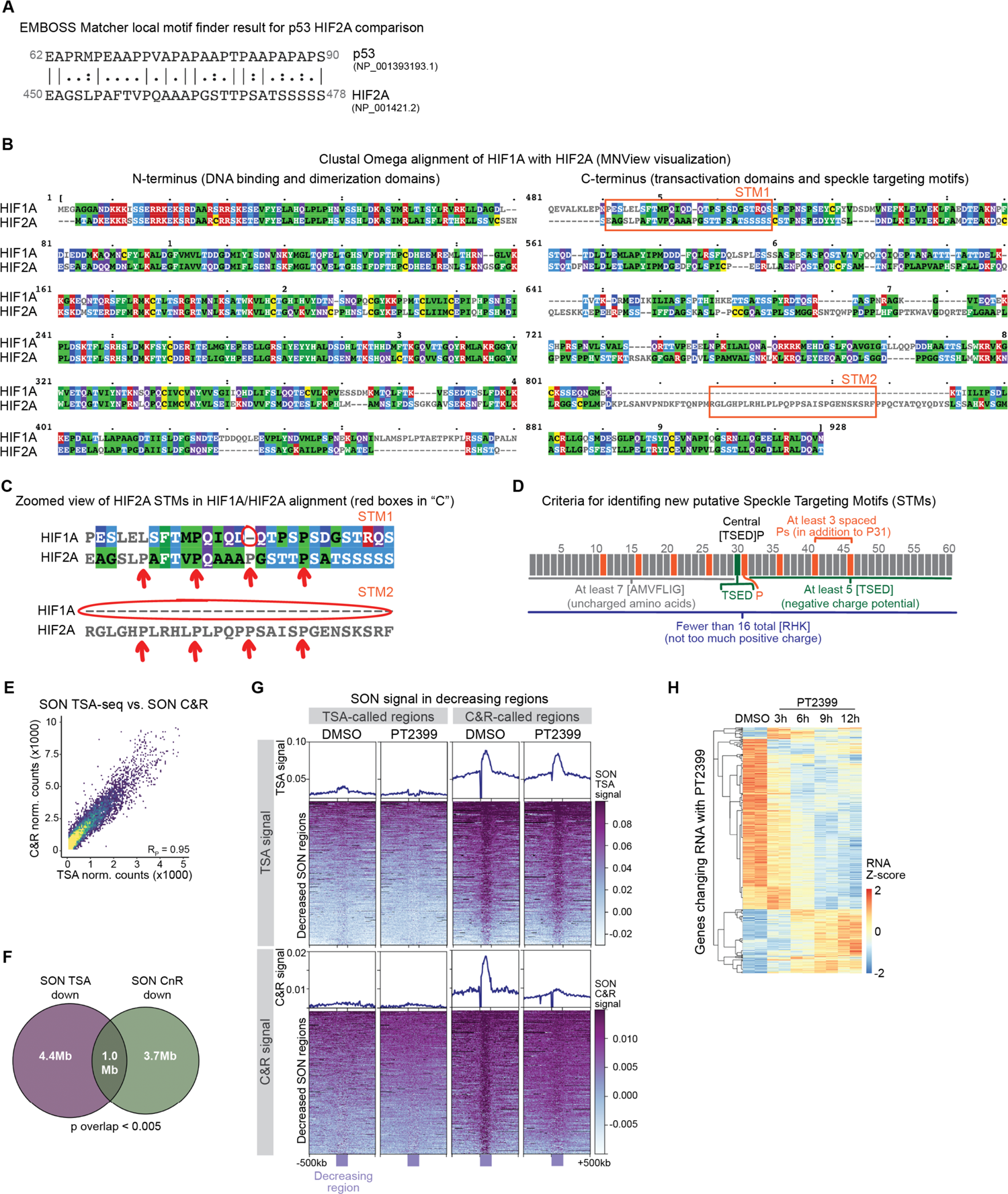
Supplement to Figure 2. A) EMBOSS Matcher alignment showing the best match local motif between p53 (amino acids 62-90) and HIF-2α (amino acids 450-478). B) Clustal Omega alignment of HIF-1α with HIF-2α with MNView visualization. HIF-2α STMs are boxed in red. C) Zoomed in view of HIF-2α STMs in HIF-1α HIF-2α alignment from **B**. D) Criteria for *de novo* identification of speckle targeting motifs (STMs). Detailed instructions can be found on Github (https://github.com/katealexander/speckleTargetingMotif.git). E) Scatterplot showing relationship between SON TSA-seq and SON Cut&Run normalized counts, averaged over two replicates. RP – Pearson’s R. F) Venn diagram showing the base-pair overlap between regions called as decreasing SON signal in SON TSA-seq (purple) or SON Cut&Run (green) upon 3 hours of PT2399 treatment. P-value represents hypergeometric p-value of overlap. G) Heatmaps and metaplots of SON TSA-seq (top) or SON Cut&Run (bottom) signal showing regions called as decreasing upon PT2399 treatment by SON TSA-seq (left) or SON Cut&Run (right). H) Heatmap of RNA expression z-scores of changing genes in 786-O cells treated with DMSO control versus a PT2399 timecourse.

**Figure S3.**
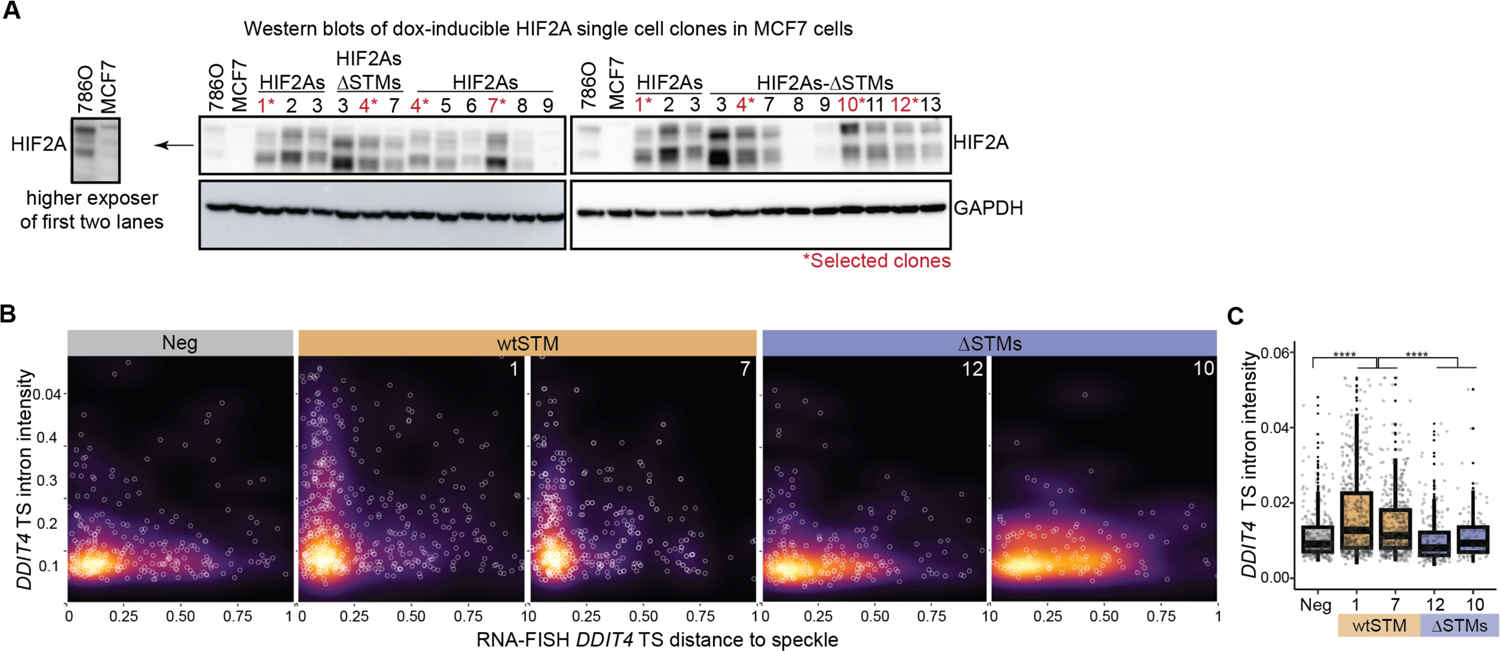
Supplement to Figure 3. A) Western blot of HIF-2α and GAPDH in 786-O cells, MCF7 cells, and MCF7 single-cell clones with dox-induced expression of HIF2As-wtSTM or HIF2As-ΔSTMs. Clones in red with asterisks denote selected clones. The same samples were loaded in the first eight lanes of both Western blots shown. B) Relationship between *DDIT4* transcription site (TS) intron intensities and distance to speckle from immunoRNA-FISH data. Each point is an individual transcription site. C) Quantification of *DDIT4* transcription site (TS) exon intensities from immunoRNA-FISH data. Each dot is an individual transcription site. Significance calculated using Wilcoxon test. **** – p < 0.0001

**Figure S4.**
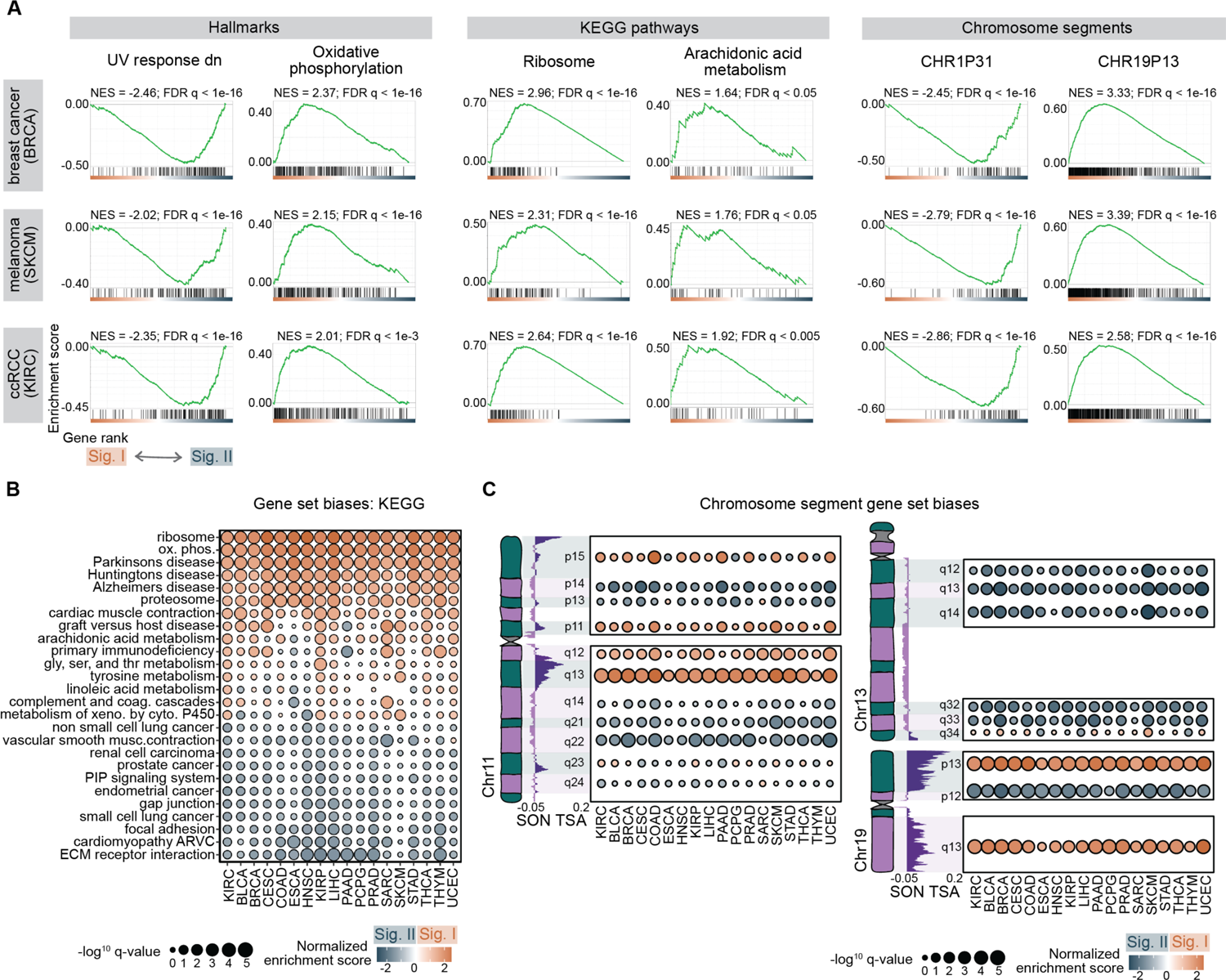
Supplement to Figure 4. A) Example gene set enrichment plots for breast cancer (BRCA), melanoma (SKCM), and ccRCC (KIRC) for Hallmark (left; see also **Fig 4A**), KEGG (middle; see also **Fig S4B**), and chromosome cytogenetic bands (right; see also **Fig 4B** and **S4C**) of gene expression biases between speckle Signature I and II patient groups. B) Gene set enrichment statistics of KEGG pathways for gene expression biases between speckle Signature I and II patient groups in ccRCC (KIRC) and other cancer types. C) Chromosome band gene set enrichment statistics for Signature I versus Signature II speckle patient groups, shown side-by-side with chromosome SON TSA-seq signal (horizontal genome browser tracks; data from Fig 2), and depictions of chromosomes 11 (left), 13 and 19 (right), with cytogenetic banding pattern shown in green and purple. See also **Fig 4B**.

**Figure S5.**
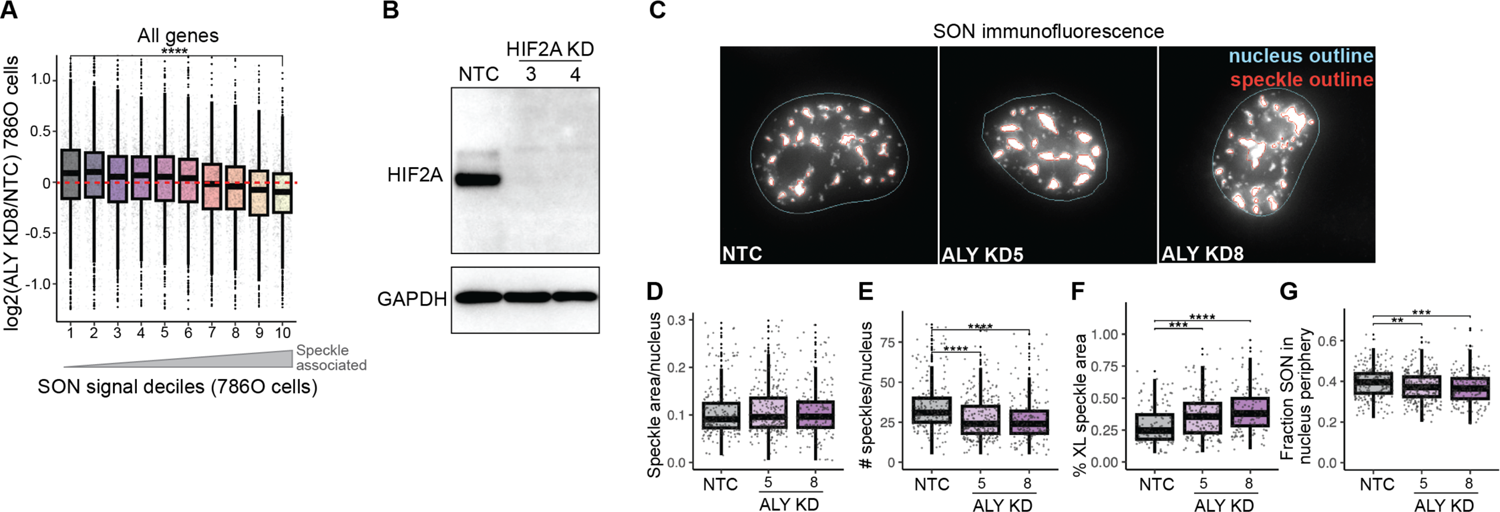
Supplement to Figure 5. A) Decile plot of the fold change upon *ALYREF* knockdown (ALY KD8; ALY KD5 in **Fig 5E**) split into deciles based on 786-O DMSO-treated SON TSA-seq signal (from Fig 2). B) Western blot of HIF-2α and GAPDH in 786-O cells treated with non-targeting control (NTC) or *HIF-2α* siRNAs (3 or 4). C) Example SON immunofluorescence images showing how speckles were called in CellProfiler (red outlines) in 786-O cells treated with NTC or *HIF-2α* siRNAs. D) Total speckle area per nucleus in 786-O cells treated with NTC or *HIF-2α* siRNAs. Each dot is an individual nucleus measurement. E) Number of speckles per nucleus in 786-O cells treated with NTC or *HIF-2α* siRNAs. Each dot is an individual nucleus measurement. F) Percentage of extra-large (XL) speckles per nucleus in 786-O cells treated with NTC or *HIF-2α* siRNAs. Each dot is an individual nucleus measurement. G) Fraction of SON signal in the nucleus periphery per nucleus in 786-O cells treated with NTC or *HIF-2α* siRNAs. Each dot is an individual nucleus measurement.

## ACKNOWLEDGEMENTS

S.L.B. acknowledges support from NIH grants RO1CA078831 and R35CA263922. K.A.A. acknowledges support from NIH grant F32CA221010 and the Marlene Shlomchik Fellowship in Cancer Research. M.C.S. acknowledges support from NIH grant R35CA220483 and DoD grant W81XWH-20-1-0856.

## AUTHOR CONTRIBUTIONS

Conceptualization: K.A.A., S.L.B., M.C.S., B.K.

Methodology: K.A.A., R.Y., S.N., E.F.J., I.P.D.

Software: K.A.A., R.Y. Validation: K.A.A., C.F., A.L.G.

Formal Analysis: K.A.A., H.H., J.L., N.B.

Investigation: K.A.A., C.F., C.L.

Resources: N.S., N.J.C., M.C.S., S.L.B., A.R., D.L.

Data Curation: K.A.A.

Writing – Original Draft: K.A.A.

Writing – Review & Editing: K.A.A., S.L.B., M.C.S., B.K. Visualization: K.A.A.

Supervision: S.L.B., M.C.S. Project Administration: K.A.A.

Funding Acquisition: S.L.B., M.C.S., K.A.A.

## METHODS

### Speckle signature and TCGA survival analysis

We took 446 protein genes annotated as “Enhanced”, “Supported”, or “Approved” for subcellular localization within nuclear speckles in The Human Protein Atlas and extracted their upper-quartile normalized RNA expression from the 30 PanCan TCGA projects that had greater than 50 samples. We performed Principal Component Analysis on these 446 speckle protein genes. In doing so, each speckle protein gene was assigned a weight (called rotation in the analysis) that was used in the analysis to separate tumor sample along the first Principal Component (PC1). The absolute value of a speckle protein gene PC1 weight thus estimates the contribution of each speckle protein gene to patient variation and the PC1 weight sign, positive or negative, reflects genes that have opposite expression patterns to one another (for example, in **Fig 1B**, “Sig-I high” and “Sig-II high” have, by definition, opposite PC1 weight signs and heatmaps show opposite expression patterns between the two groups). To compare speckle protein gene expression contributions to patient variation between cancer types, we took the pairwise Pearson’s correlation coefficients of the speckle protein PC1 weights (an example single pairwise comparison is shown in **Fig S1A**, right scatterplot; the correlation coefficients of each cancer-to-cancer pairwise comparison are summarized in **Fig 1Aiii** heatmap).

To obtain a set of speckle protein genes that consistently contributed to patient variation in many cancer types, we flipped the rotation signs so that the speckle protein gene, SON, was always assigned a negative weight.

We extracted the speckle protein genes that had consistently signed rotations across 22 cancer types (the 22 cancer types in **Fig 1Aiii** at the bottom of the heatmap that showed highly similar speckle protein gene PC1 weights to one another) and used these 117 speckle protein genes to calculate speckle scores (40 Signature I-high speckle protein genes; 77 Signature II-high speckle protein genes). Speckle scores were calculated by taking the z-scores of speckle protein gene expression, calculated per cancer, and applying the following formula: sum((z-score Sig I speckle protein gene)*1/(number Sig I speckle protein genes)) + sum((z-score Sig II speckle protein gene)*-1/(number Sig II speckle protein genes). In this manner a speckle score was assigned to sample so that it would be strongly positive for tumors with the strongest Signature I expression pattern and strongly negative for tumors with the strongest Signature II expression pattern. Speckle score was then used to separate samples into groups for Kaplan Meier and gene expression analysis between the two groups. For detailed instructions, scripts, and output files for the RNA-based speckle signature, see the speckle signature GitHub page: https://github.com/katealexander/speckleSignature.git.

### Speckle targeting motif

To identify speckle targeting motifs across the proteome, we first used Motif Search to download all the motifs in the human proteome that followed this rule: x(30)-[TSED]-P-x(30), meaning any 30 amino acids followed by a threonine, serine, glutamine, or asparagine, followed by a proline, followed by any 30 amino acids. From these motifs, we applied the following rules to identify speckle targeting motifs: 1) no more than 4 prolines in a row, 2) at least 3 spaced prolines of 7 possibilities spaced every 5 amino acids from the central [TSED]P (Python indexes 11, 16, 21, 26, 36, 41, 46), 3) at least 5 of TSED on the C-terminal side of the central [TSED]P, 4) at least 7 of AMVFLIG on the N-terminal side of the central [TSED]P, 5) fewer than 16 total RHK amino acids (summarized in **Fig S2D**). We note that these rules are for the purpose of identifying new putative speckle targeting motifs and may be modified as additional information about the required biochemistries of speckle targeting becomes available in future studies. Detailed instructions can be found on the speckle targeting motif GitHub page: https://github.com/katealexander/speckleTargetingMotif.git. Lists of the identified STM-containing proteins (**Table S2**), and their STM sequences (**Table S3**) are included in Supplementary Materials.

### Tissue culture

786-O cells were grown in RPMI media with 10% FBS under atmospheric oxygen and 5% carbon dioxide. For HIF-2α inhibition experiments, 786-O cells were treated with 2µM of PT2399 or equivalent volume of DMSO vehicle control for 3 hours. MCF7 cells were grown in DMEM media with 10% FBS under atmospheric oxygen and 5% carbon dioxide. To generate MCF7 cells with dox-inducible HIF-2α, we packaged the reverse tetracycline controlled transactivator plasmid (Lenti CMV rtTA3 Blast [w756-1]) into viruses, infected MCF7 cells, and performed selection for cells with stably-integrated rtTA using blasticidin. MCF7-rtTA cells were then infected with viruses containing stabilized HIF-2α with wild type STMs or with deleted STMs under the control of a TRE3G promoter (pRetroX-TRE3G HIF-2α) and selected for stable integration using puromycin. After puromycin selection, single cell cloning was performed by diluting cells, plating in a 96 well plate and selecting wells with single clones. Individual clones were screened for HIF-2α levels by Western blotting. For virus generation, HEK293T cells were transfected with pPAX2, pVSV-G, and the lentiviral vector (rtTA or pRetroX-TRE3G) using Lipofectamine2000, as described^50^, and infected MCF7 cells with a 1:1 ratio of virus-containing media and MCF7 growth media. To induce HIF-2α expression, doxycycline was dissolved in water to a 30mM stock solution and added to MCF7 cells at a final concentration of 3µM.

### HIF-2α ChIP-seq

786-O cells treated with DMSO vehicle control or 2µM PT2399 for 3 hours were crosslinked in formaldehyde (1% final concentration) for 10 minutes. Crosslinked cells were quenched with glycine (125mM final) for 5 minutes, followed by two washes in ice-cold PBS. Nuclei were isolated from approximately 20 million cells as previously described^51^, and chromatin was sheared using a Covaris S220. Immunoprecipitation was performed using 500µg of sheared chromatin lysate and 5µg of HIF-2α antibody (Novus NB100-122) pre-conjugated to protein G beads (Invitrogen). Lysate and antibody-beads were incubated for 16 hours at 4°C with rotation and then washed four times in wash buffer [50mM HEPES-HCl(pH 8), 100mM NaCl, 1mM EDTA, 0,5mM EGTA, 0.1% sodium deoxycholate, and 0.5% N-laurylsarcosine], followed by one wash in ChIP final wash buffer [1x tris-EDTA (TE) Buffer and 50mM NaCl]. Immunoprecipitated DNA was eluted from washed beads, reverse cross-linked overnight and purified. The resulting DNA fragments were prepared for sequencing using the NEBNext Ultra II DNA Library Prep Kit for Illumina. Library sizes were determined on a Bioanalyzer, and concentration determined using NEBNext Lib Quant Kit (E7630, NEB). Input and pulldown libraries were sequenced with an Illumina NextSeq550 with 42bp per read using the NextSeq 500/550 High Output 75-cycle v2.5 kit.

### SON TSA-seq

TSA-seq was performed on 786-O cells treated with DMSO or PT2399 for 3 hours using an antibody against SON (ab121759). SON TSA-seq was carried out using the protocol for SON TSA-seq 2.0 on attached cells with Condition E as described (as in ^19^; labeling with 50% sucrose, 1:300 tyramide-biotin, and 0.0015% hydrogen peroxide for 30 minutes), with the following minor modifications: DNA was fragmented to an average size of 250bp using a Covaris S220 and the resulting DNA fragments were prepared for sequencing using the NEBNext Ultra II DNA Library Prep Kit for Illumina. Library sizes were determined on a Bioanalyzer, and concentration determined using NEBNext Lib Quant Kit (E7630, NEB). Libraries were sequenced with an Illumina NextSeq550 with 42bp per read (total of 84bp) using the NextSeq 500/550 High Output 75-cycle v2.5 kit.

### SON Cut&Run

Cut&Run was performed similarly to previously described^35^ on 786-O cells treated with DMSO or PT2399 for 3 hours. For each condition, nuclei from ∼600k cells was isolated using nuclear isolation buffer [10mM HEPES-KOH (pH 7.9), 10mM KCl, 0.1% NP40, 0.5mM spermidine, 1x Halt protease inhibitor cocktail] and bound to concanavalin A lectin beads which had been washed in binding buffer [BioMag Plus; binding buffer: 20mM HEPES-KOH (pH 7.9), 10mM KCl, 1mM CaCl2, 1mMnCl2], as described. After bead binding, samples were split into two tubes, one for each antibody, where binding buffer was replaced with 1:100 antibody diluted in blocking buffer [20mM HEPES-KOH (pH 7.5), 150mM NaCl, 0.1% BSA, 0.5mM spermidine, 1x Halt protease inhibitor cocktail, 2mM EDTA](IgG antibody: Millipore 06-371; SON antibody: abcam ab121759) and incubated overnight at 4°C. Samples were washed in washing buffer [20mM HEPES-KOH (pH 7.5), 150mM NaCl, 0.1% BSA, 0.5mM spermidine, 1x Halt protease inhibitor cocktail], and treated with pA-MNase and CaCl2 for 30 minutes on an ice-cold pre-chilled metal block on ice. The reaction was stopped by addition of STOP buffer [200mM NaCl, 20mM EDTA, 4mM EGTA, 50µg/mL RNase A, 40µg/mL glycogen] while gently vortexing, and DNA fragments were released by incubation at 37°C for 10 minutes. The DNA fragments were collected from bead supernatants, treated with Proteinase K for 10 minutes at 70°C, then subjected to PCI-choroform purification. Libraries from DNA fragments were made using the NEBNext Ultra II DNA Library Prep Kit for Illumina. Library sizes were determined on a Bioanalyzer, and concentration determined using NEBNext Lib Quant Kit (E7630, NEB). Libraries were sequenced with an Illumina NextSeq550 with 42bp per read (total of 84bp) using the NextSeq 500/550 High Output 75-cycle v2.5 kit.

### SON TSA-seq and SON Cut&Run analysis

SON TSA-seq or SON Cut&Run reads were aligned to the human reference genome assembly GRC37/hg19 using Bowtie2 allowing for a maximum fragment size of 1000 base pairs^52^. PCR duplicates were removed using Picard, and SON signal was quantified over sliding windows of 50 or 100kb slid by 1/10^th^ of the window size using DiffBind with input (TSA-seq) or IgG (Cut&Run) subtraction^53^. Differential 50kb and 100kb windows identified from DiffBind were merged using BEDTools^54^, resulting in differential domains. Genes within differential domains were then extracted using Python. Genes within decreasing domains that had an adjusted p-value of less than 0.01 in either SON TSA-seq or SON Cut&Run were considered to have speckle association regulated by HIF-2α; genes were considered unchanging if they were in domains with an adjusted p-value of greater than 0.1 by both methods. For more detailed analysis instructions and associated scripts, see Github: https://github.com/katealexander/TSAseq-Alexander2020/tree/master/genomicBins_DiffBind.

### immunoDNA-FISH

Oligopaint DNA-FISH probes for genes were designed across a 50kb region centered on the transcription start site as described^55^. ImmunoDNA-FISH was performed first with the standard immunofluorescence protocol, followed by 10 minutes in 4% paraformaldehyde to fix the secondary antibody prior to DNA-FISH. Specifically, cells were fixed in 4% paraformaldehyde for 10 minutes, permeabilized using 0.1% Triton X-100 in PBS for 10 minutes, incubated with 1:200 SON antibody diluted in PBS overnight at 4°C, and incubated in 1:200 anti-rabbit A488 secondary for 1 hour at room temperature, with 3×5 minute washes in PBS after fixation and antibody steps. After fixation and PBS washes, DNA-FISH was performed as described^56^.

### immunoRNA-FISH

RNA-FISH probes were designed using Stellaris probe design software, inputting *DDIT4* exons (for exonic probe set) and introns (for intronic probe set). Immunofluorescence was performed as in immunoDNA-FISH, with the exception that primary and secondary antibodies were used at a 1:1000 dilution. After immunofluorescence, cells were incubated overnight in 70% ethanol at 4°C. RNA-FISH was then performed as previously described^57^. *DDIT4* exonic probes were labelled with Quasar 570, *DDIT4* intronic probes were labelled with CAL Fluor Red 610, and a mouse anti-rabbit A488-labelled secondary was used.

### RNA-seq

For RNA-seq of 786-O cells, cells were treated with DMSO for 12 hours or with 2µM of PT2399 for 3, 6, 9, or 12 hours. Cells were lysed in TRIzol (15596018, Thermo Fisher Scientific) and snap frozen. RNA was then isolated using chloroform extraction, followed by QIAGEN RNeasy Mini Kit isolation (#74106), including DNA digestion with DNase. Poly(A) + RNA was then isolated using double selection with poly-dT bead (E7490, NEB), and RNA-seq libraries were prepared using a NEB-Next Ultra II Directional Library Prep Kit for Illumina (E7760, NEB). Library sizes were determined on a Bioanalyzer, and concentrations determined using NEBNext Lib Quant Kit (E7630, NEB). Libraries were sequenced on an Illumina NextSeq 550, using paired-end sequencing of 42 bases per read. For RNA-seq of ccRCC and normal adjacent tissues, the above protocol was applied to flash frozen tissue samples with the added step of manually homogenizing the tissue during the TRIzol step prior to chloroform extraction.

### Definitions of HIF-2α target genes

Because exact HIF-2α target genes are known to be context dependent^58^, our working definition of what constitutes a HIF-2α target gene varied depending on the analysis/cell type. For 786-O cells (**Fig 2H-K**), 874 genes decreasing with PT2399 at an adjusted p-value of less than 0.1 at 3, 6, 9, or 12 hours were used. For MCF7 cells (**Fig 3H-I** and **Fig 4H-I**), 1169 genes increasing with dox-inducible HIF2As-wtSTM or HIF2As-ΔSTMs at an adjusted p-value of less than 0.1 were used. For analysis of TCGA tumor samples (**Fig 4D-G**), we combined the 786-O HIF-2α targets, MCF7 HIF-2α targets, and a list of published HIF-2α targets^59^ and applied the additional filter of having increased expression in either tumor patient group compared to tissue-adjacent normal samples (p-value < 0.05, log2 fold change > 0.3), for 986 genes.

### Molecular cloning

pVHL-stabilized HIF-2α (P405A/P531A) was amplified by PCR from the HA-HIF2alpha-P405A/P531A-pcDNA3 plasmid (Addgene, 18956) using primers BamHI-HIF2Af and AscI-HIF2Ar using NEB Q5 High-Fidelity 2X Master Mix according to the instructions of the manufacturer. The resultant PCR product was purified using Qiagen MinElute PCR Purification Kit. pRetroX-Tre3G plasmid and purified PCR product were digested with the BamHI and AscI restriction enzymes, gel purified using Qiagen QIAquick Gel Extraction Kit, ligated using T4 DNA ligase, and transformed into Stbl3 chemically competent *E. coli*. Individual colonies were picked, miniprepped, and correct pRetroX-Tre3G-HIF2A-P405A/P531A clones were confirmed by Sanger sequencing. STM mutations were introduced into pRetroX-Tre3G-HIF2A-P405A/P531A sequentially using mutagenesis PCR with first the STM1Af and STM1Ar primers, then upon successful generation of the first STM deletion, applying mutagenesis PCR with the second STM2Af and STM2Ar primers. Mutagenesis PCR was performed using NEB Q5 High-Fidelity 2X Master Mix according to the instructions of the manufacturer. The resulting PCR product was treated with kinase, ligase, and DpnI (KLD enzyme mix; NEB M0554) and transformed in to Stbl3 chemically competent *E. coli*. Correct clones were confirmed with Sanger sequencing.

### Western blotting

Cells were lysed in RIPA buffer containing 1% NP-40, 150mM NaCl, 50mM TRIS-Cl (pH 8.0), and 1% SDS supplemented with cOmplete, EDTA-free protease inhibitor (11873580001, Roche) for 15 minutes at 4°C in a Bioruptor. Cell lysates were centrifuged at max speed at 4°C for 10 minutes, and the membrane pellets were discarded. Protein concentration was determined by bicinchroninic acid assay (BCA) protein assay (#23227, Life Technologies), after which equal amounts of protein were loaded and separated by polyacrylamide gel electrophoresis. The following antibodies were used: HIF-2α (NB100-122, Novus) and GAPDH (10R-G109a, Fitzgerald Industries).

### Gene ontology analysis

For Gene Ontology (GO) analysis of HIF-2α target gene subgroups (**Fig 2K** and **Fig 4G**), we first extracted GO Biological Process terms using the R package clusterProfiler function enrichGO for all HIF-2α target genes with p and q-value cutoffs of 0.01. We then extracted the enrichment statistics for each subset of HIF-2α target genes and plotted the terms that were differential between groups. For simplicity in representation, we displayed the most significant GO term for terms that were similar to one another, (e.g. “protein localization to nucleus” and “positive regulation of protein localization to nucleus” or “cell response to interleukin-4” and “response to intereukin-4”). For full instructions, scripts, and gene lists used see Github: https://github.com/katealexander/geneOntologyOfSubsets.git.

### Gene set enrichment analysis

Gene set enrichment analysis (GSEA) was performed using gsea_4.3.2 for each cancer type using the Hallmark (H), Positional (C1), and KEGG (C2:CP:KEGG) gene sets. GSEA enrichment scores and -log of FDR q-values were visually represented in bubble plots using R as described in Github: https://github.com/katealexander/makeBubblePlotFromGSEAstats.git. Included in this Github repository are bubble plots of the GSEA results for each chromosome arm, in the folder “chromosomeEnrichments”, as well as the input files used for GSEA analysis.

### Integrating speckle signature, expression, and speckle association datasets

Per-gene SON signal from 786-O SON TSA-seq DMSO data was calculated as previously described^21^. Then, to integrate data between datasets, two types of approaches were used: 1) splitting one dataset into n-tiles and graphing the values of the other dataset or 2) splitting one dataset by differential gene expression and graphing the values of the other dataset. For detailed instructions and scripts used, see Github: https://github.com/katealexander/dataIntegration-Alexander2023.git.

### Speckle protein-protein interactome network

The speckle protein-protein interactome network to identify speckle-resident functional complexes was generated in StringDB with the following filters: physical subnetwork, high confidence, experiments and datasets, 200 interactors in first shell, 100 interactors in second shell. Enriched non-overlapping functional categories (CL:1598, GO:0006325, GO:0071005, GO:0071007, GO:0071006, GO:1903311, GO:0006281, and GO:0006366) were selected for visualization and analysis of speckle signature biases. Detailed instructions, files, and scripts used are available at Github: https://github.com/katealexander/speckleProteinInteractome.git.

### Knockdowns

HIF-2α or ALYREF were knocked down in 786-O cells using FlexiTube siRNAs (Qiagen) and compared to a non-targeting siRNA control (Qiagen Allstars Neg Ctrl). 786-O cells were split one day prior to transfection and plated at 50-60% confluency. siRNAs were transfected into cells using the DharmaFECT 1 Transfection Reagent (Horizon T-2001-03) according to the instructions of the manufacturer. siRNA-treated 786-O cells were split into fresh media 24 hours after transfection and harvested 48 hours after transfection. Knockdown efficiency was determined by Western blot (HIF-2α knockdown) or immunofluorescence (ALYREF knockdown).

### Immunofluorescence in tissue culture cells

Immunofluorescence of SON (Abcam ab121759, used at 1:1000) and ALYREF (Santa Cruz sc-32311, used at 1:200) was performed in 786-O cells with *ALYREF* knockdown. 786-O cells treated with non-targeting control or *ALYREF* siRNAs were plated onto 8-well glass coverslip plates (LabTek 155411), harvested by fixation for 10 minutes with 4% PFA (Chem Cruz sc-281692), washed 3×5 minutes in PBS, permeabilized for 10 minutes in 0.1% Triton X-100 diluted in PBS, incubated in primary antibody diluted in PBS overnight at 4°C, washed 3×5 minutes in PBS, incubated in secondary antibody (Invitrogen Alexa Fluor 488 goat anti-rabbit IgG at 1:1000; Alexa Fluor 647 goat anti-mouse IgG at 1:200) for 1 hour at room temperature, washed for 5 minutes in PBS, incubated in DAPI (Invitrogen D1306; 1:50,000 of 5mg/mL) for 10 minutes at room temperature, washed 3×5 minutes in PBS, and submerged in mounting media (20mM Tris pH 8.0, 0.5% N-propyl gallate, 90% glycerol).

### Analysis of knockdown efficiency from immunofluorescence data

Knockdown of ALYREF was quantified in immunofluorescence 20x imaging data by taking the median nuclear intensity of ALYREF, using CellProfiler to perform maximum projections, identify nuclei, and quantify intensities. Full instructions, example data, the CellProfiler pipeline used on maximum projections, and Python and R scripts used for post-processing can be found at GitHub: https://github.com/katealexander/cellProfilerPipelines/tree/main/knockdownQuantification.

### Analysis of speckle phenotypes from 60x immunofluorescence data

Speckle phenotypes from 60x imaging data were determined on maximum projections of z-stacks using CellProfiler to perform maximum projections, identify nuclei, identify speckles, and calculate intensity measurements of the SON speckle marker. Full instructions, example data, the CellProfiler pipeline used on maximum projections, and Python and R scripts used for post-processing can be found at GitHub: https://github.com/katealexander/cellProfilerPipelines/tree/main/specklePhenotypes60x.

### SON immunofluorescence in FFPE tissue sections

A tissue array of formalin-fixed paraffin-embedded 5µm ccRCC with matching tumor adjacent tissue sections and associated survival and tumor grade data was obtained from USbiomax (HKID-CRC180SUR-01). The slide was baked for 2 hours at 60°C to prevent tissue detachment, deparaffinized using 3×5 minute washes in Xylenes, and rehydrated in a series of 2×10 minute incubations in each of 100%, 95%, 80%, 70%, and 50% EtOH followed by 2×5 minute washes in each of deionized water, TBS (10mM Tris, 150mM NaCl, pH7.5), and deionized water. Antigen retrieval was accomplished by a 5-minute pressure cook of the slide submerged in 1X HIER buffer (abcam ab208572). Deionized water was slowly added to the pressure-cooked slide in HIER buffer, doubling the total volume to gradually cool the slide, then the slide was removed and incubated 2×5 minutes in deionized water. Following antigen retrieval, samples were blocked in 10% goat serum diluted in PBS with 0.2% Triton X-100 for 1.5 hours at room temperature. Primary SON antibody (abcam ab121759, used at 1:100) diluted in 1% goat serum with 0.2% Triton X-100 in PBS was applied to the samples, then samples were covered with a flexible hybridization cover (Invitrogen H18202) and incubated in a humidified chamber overnight at 4°C. Samples were washed 2×10 minutes in 1% goat serum with 0.2% Triton X-100 diluted in PBS, incubated in secondary antibody (Invitrogen Alexa Fluor 647 goat anti-rabbit IgG at 1:100) for 2 hours at room temperature, washed 1×10 minutes in 1% goat serum with 0.2% Triton X-100 diluted in PBS, DAPI-treated (Invitrogen D1306; 1:50,000 of 5mg/mL) for 10 minutes in 1% goat serum with 0.2% Triton X-100 diluted in PBS, washed 2×10 minutes in 1% goat serum with 0.2% Triton X-100 diluted in PBS, and mounted in mounting media (20mM Tris pH 8.0, 0.5% N-propyl gallate, 90% glycerol) with a coverslip.

### SON immunofluorescence FFPE 20x image analysis: intensity measurements, survival, tumor grade comparison

After imaging the entirety of each tissue sample at 20x, maximum projections of z-stacked images were performed, nuclei were called, and intensity measurements were calculated in CellProfiler. For each sample, median nuclei measurements were calculated, and survival analysis was performed by splitting the cohort by the top and bottom 50% of each measurement. Likewise, median measurements were used to compare ccRCC tumor to normal adjacent tissue and to assess measurement differences based on tumor grade. Full instructions, processed data, Kaplan Meier plots for each measurement, violin plots of tumor versus normal for each measurement, the CellProfiler pipeline, and Python and R scripts used for processing can be found at GitHub: https://github.com/katealexander/cellProfilerPipelines/tree/main/specklePhenotypes20xWithSurvivalAnalysis.

### ccRCC and normal primary renal cortex sample collection and processing

De-identified ccRCC and tumor adjacent normal primary renal cortex samples were collected directly at the Hospital of the University of Pennsylvania, Division of Hematology and Oncology. Patients provided written consent allowing the use of discarded surgical samples for research purposes on an Institutional Review Board approved protocol (IRB 843862). Individuals were excluded if they were known to be positive for HBV, HCV, or HIV. Tissues were collected in sterile conditions and transported on ice to the laboratory for further processing. Handling of samples was performed in accordance with Biosafety Level 2 standards with sterile material.

Processing consisted of transferring the patient sample into a sterile petri dish with PBS and cutting fragments for patient-derived xenograft (PDX) implantation (5mm^3^), long-term preservation (5mm^3^) at −80°C in 10% DMSO in fetal bovine serum (FBS), and flash freezing in liquid nitrogen for molecular studies (2-5mm^3^).

Patient-derived xenograft (PDX) engraftment was performed within the Stem Cell and Xenograft Core (RRID:SCR_010035) at the University of Pennsylvania and in accordance with the Institutional Animal Care and Use Committee (IACUC protocol #806944). Tumor fragments were subcutaneously implanted into 4- to 6-week-old NSG recipient mice (Jackson Laboratories, Strain #:005557) with Matrigel (Corning #354234). Tumor growth was monitored weekly. Animals were sacrificed and tumors harvested similarly to the human sample protocol above once tumors reached the maximum allowed size (2000mm^3^).

For RNA-seq and imaging analysis, individual frozen samples preserved in liquid nitrogen were retrieved and sliced into two pieces, with one piece for formalin fixation and paraffin embedding followed by immunofluorescence as described above, and the second piece for RNA-seq as described above. For formalin fixation and paraffin embedding, tissue samples were fixed in 4% PFA (Chem Cruz sc-281692) for 48 hours at 4°C, washed 3x in PBS, incubated for 1 hour in 50% EtOH, stored in 70% EtOH, then paraffin-embedded and sectioned by The Molecular Pathology and Imaging Core (MPIC) at the University of Pennsylvania. H&E staining was performed by the MPIC, and the identity of the tissue sample (ccRCC tumor or normal renal cortex) was confirmed by a pathologist.

### Microscopy

Microscopy was performed on a Nikon Ti2E Motorized XYZ Inverted Fluorescence Microscope with Perfect Focus System using a 20x Objective Lens (N.A. 0.75, W.D. 1.0mm, F.O.V. 25mm) or a 60x Oil Immersion Objective Lens (N.A. 1.4, W.D. 0.13mm, F.O.V. 25mm) with an LED fluorescent light source and the following filter sets: C-FL DAPI SOLA Hard Coat (Nikon 96359), C-FL YFP Hard Coat (Nikon 96363), 49304 ET Gold Filter Set (Nikon 77074178), 49311 ET – Red #3 Narrow-band FISH Filter Set (Nikon 77074733), and 49307 ET Far Red FISH Filter Set (Nikon 77074477). For 20x imaging, a set of 7-9 z-stacks spaced 1.3µm apart were imaged spanning the entire focal plane of the imaging sample. For 60x imaging, a set of 29-35 z-stacks spaced 0.29µm apart were imaged spanning the entire focal plane of the imaging sample.

